# Relationship between Dynamic Instability of Individual Microtubules and Flux of Subunits into and out of Polymer

**DOI:** 10.1101/609701

**Authors:** Ava J. Mauro, Erin M. Jonasson, Holly V. Goodson

## Abstract

Behaviors of dynamic polymers such as microtubules and actin are frequently assessed at one or both of two scales: (i) net assembly or disassembly of bulk polymer, (ii) growth and shortening of individual filaments. Previous work has derived various forms of an equation to relate the rate of change in bulk polymer mass (i.e., flux of subunits into and out of polymer, often abbreviated as “*J*”) to individual filament behaviors. However, these versions of this “*J* equation” differ in the variables used to quantify individual filament behavior, which correspond to different experimental approaches. For example, some variants of the *J* equation use dynamic instability parameters, obtained by following particular individuals for long periods of time. Another form of the equation uses measurements from many individuals followed over short time steps. We use a combination of derivations and computer simulations that mimic experiments to (i) relate the various forms of the *J* equation to each other; (ii) determine conditions under which these *J* equation forms are and are not equivalent; and (iii) identify aspects of the measurements that can affect the accuracy of each form of the *J* equation. Improved understanding of the *J* equation and its connections to experimentally measurable quantities will contribute to efforts to build a multi-scale understanding of steady-state polymer behavior.

## 1 INTRODUCTION

Behaviors of dynamic polymers such as microtubules (MTs) and actin are frequently assessed at the scales of populations and/or individual filaments. Previous work has investigated various forms of an equation that use quantities describing individual filament dynamics to estimate the rate of change in the population’s polymer mass (e.g., (Hill & Chen, 1984; Walker et al., 1988; Verde, Dogterom, Stelzer, Karsenti, & Leibler, 1992; Dogterom & Leibler, 1993; Komarova, Vorobjev, & Borisy, 2002)). The rate of change in the population’s polymer mass is also described as the flux (abbreviated as *J*) of subunits into and out of polymer, so we refer to this equation as the *J* equation. The versions of the *J* equation differ in the particular variables used to quantify individual filament behavior, which correspond to different experimental approaches (e.g., following particular individuals for long times (Walker et al., 1988), or many individuals each over short time steps (Komarova et al., 2002)). In this paper, we relate the various forms of the *J* equation to each other and use computational simulations to demonstrate these relationships. We also discuss aspects of the measurements that can affect the accuracy of the output of each form of the *J* equation. This paper focuses on microtubules, but should apply to steady-state polymers more broadly. For definitions of abbreviations and terms used in this paper, please see Table 1.

**Table 1:** Definitions of terms and abbreviations.

| Term or Abbreviation | Definition |
| --- | --- |
| MT | Microtubule |
| [free tubulin] | Concentration of tubulin dimers (subunits) in solution |
| [polymerized tubulin] | Concentration of tubulin dimers (subunits) in polymerized form |
| [total tubulin] | Total concentration of tubulin dimers (subunits) in a system<br>= [free tubulin] + [polymerized tubulin] |
| DI | Dynamic instability: behavior in which MTs stochastically switch between periods of extended growth and shortening |
| DI parameters | Four measurements ($V_g$, $V_s$, $F_{cat}$, and $F_{res}$ as defined below) commonly used to quantify DI behavior |
| $V_g$ | Growth velocity of individual MTs during growth phases |
| $V_s$ | Shortening velocity of individual MTs during shortening phases |
| $F_{cat}$ | Frequency of catastrophe = number of catastrophes / time in growth |
| $F_{res}$ | Frequency of rescue = number of rescues / time in shortening |
| $J$ | The rate of change in average MT length = drift coefficient of MT ends = per MT average flux of tubulin into and out of polymer (e.g., in $\mu\text{m/s}$);<br>or the rate of change in [polymerized tubulin] = flux of tubulin into and out of a population's polymer mass (e.g., in $\mu\text{M/s}$) |
| CC | Critical concentration |
| $CC_{NetAssembly}$ | CC above which the steady state value of $J$ is positive ($J > 0$) (e.g., (Walker et al., 1988; Dogterom & Leibler, 1993; Jonasson et al., 2019)). At [free tubulin] above $CC_{NetAssembly}$, the average MT length will increase and individual MTs will experience net growth over time. |
| $CC_{Elongation}$ | CC above which $V_g$ is positive ($V_g > 0$), as determined by linear extrapolation to $V_g = 0$ from a plot of $V_g$ versus [free tubulin] (e.g., (Walker et al., 1988; Jonasson et al., 2019)). At [free tubulin] above $CC_{Elongation}$, individual MTs can exhibit extended but transient growth phases. |
| Competing simulation | Simulation of a closed system where [total tubulin] is held constant for the duration of the simulation, and MTs compete for tubulin |
| Constant [free tubulin] simulation | Simulation of an open (non-competing) system where [free tubulin] is held constant for the duration of the simulation |
| Dilution simulation | Simulation in which MTs are grown under competing conditions at very high [total tubulin] (ideally, [total tubulin] $\gg CC_{NetAssembly}$) to produce long MTs, which are then transferred into non-competing conditions at various values of [free tubulin] |
| Polymer-mass steady state | A steady state in which [polymerized tubulin] (or, alternatively, average MT length) has reached a value that is constant with time ($J = 0$, "bounded growth" regime in (Dogterom & Leibler, 1993; Fygenson et al., 1994)). This steady state occurs in competing systems at any [total tubulin] and in non-competing systems at [free tubulin] $< CC_{NetAssembly}$ (e.g., (Jonasson et al., 2019)). |
| Polymer-growth steady state | A steady state in which [polymerized tubulin] (or, alternatively, average MT length) increases at a constant rate ($J > 0$, "unbounded growth" regime in (Dogterom & Leibler, 1993; Fygenson et al., 1994)). This steady state occurs in non-competing systems at [free tubulin] $> CC_{NetAssembly}$ (e.g., (Jonasson et al., 2019)). |

### 1.1 Flux of subunits into and out of polymer

The flux (*J*) of subunits into and out of polymer has been used to quantify behaviors of polymers such as microtubules and actin (e.g., (Carlier, Pantaloni, & Korn, 1984b; Carlier, Hill, & Chen, 1984a; Verde et al., 1992; Vavylonis, Yang, & O’Shaughnessy, 2005). In a traditional flux measurement experiment, the relationship between flux and subunit concentration is determined by first growing filaments to long lengths in an environment with a high concentration of subunits, then “diluting” (transferring) samples into known concentrations of subunits and assessing the rate at which polymer assembles or disassembles (e.g., (Carlier et al., 1984a; Carlier et al., 1984b). Figure 1A,D shows schematic representations of *J* as a function of subunit concentration as obtained from such a dilution experiment (panel **A**) and from another type of experiment where [free subunit] is held constant for the entire duration of the experiment (panel **D**).

**Figure 1:**
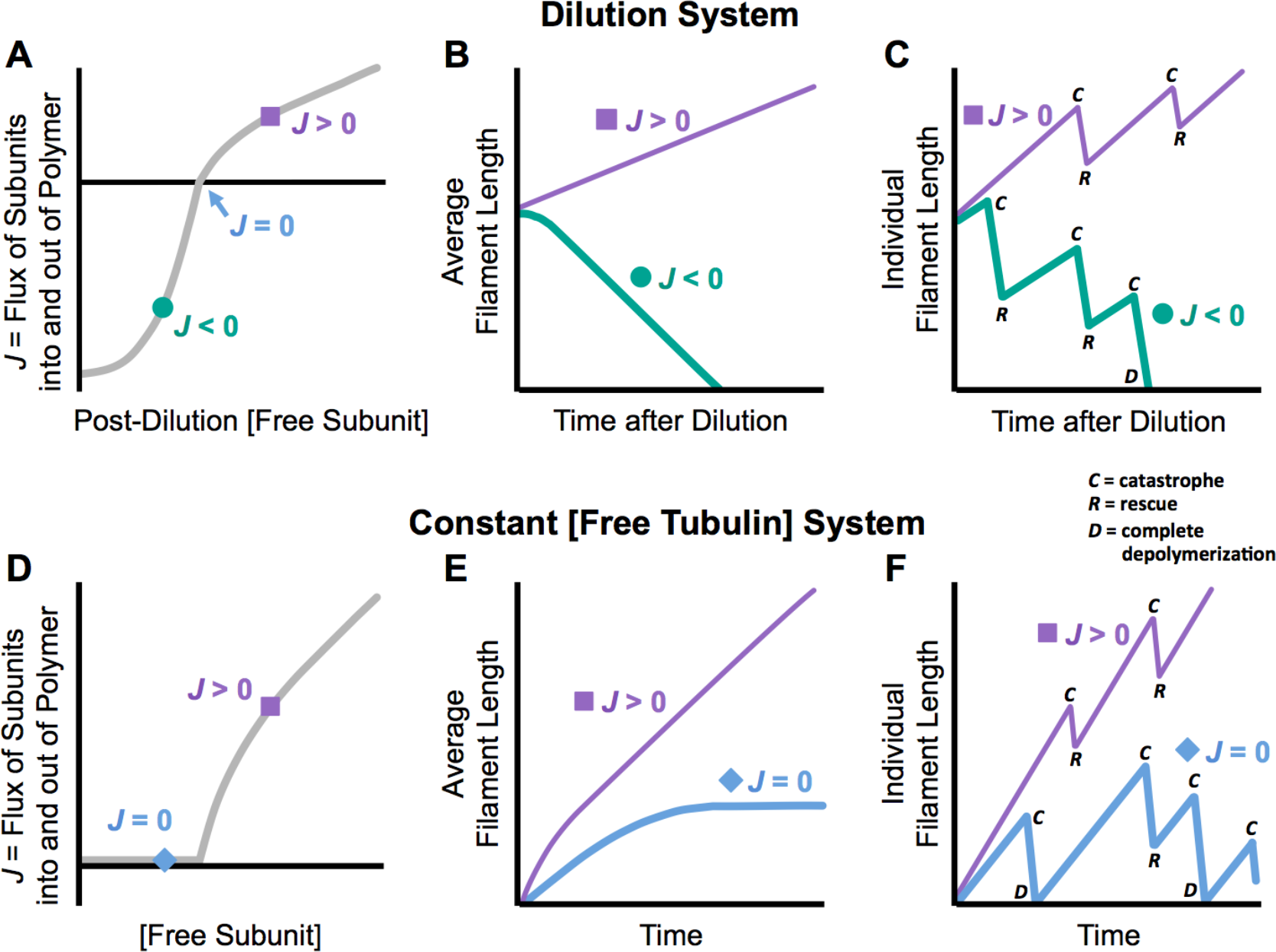
Schematic representation of flux and dynamic instability in dilution experiments and constant [free subunit] experiments. In dilution experiments (top row), polymer is grown under competing conditions at high [total subunit] to produce long filaments and then “diluted” into various values of [free subunit] under non-competing conditions (e.g., (Carlier et al., 1984a). In constant [free subunit] experiments (bottom row), the value of [free subunit] is held constant for the duration of the experiment. See also the descriptions of competing, constant [free subunit], and dilution experiments in Table 1. **(*A,D*)** Flux of subunits into and out of polymer as a function of [free subunit]. **(*B,E*)** Average filament length of a population versus time. Each curve in panels **B,E** corresponds to a single value of [free subunit] in panels **A,D**, as indicated by the symbol shapes. **(*C,F*)** Representative individual filament length versus time. Each length history in panels **C,F** represents an individual from the populations in panels **B,E** at the corresponding single value of [free subunit] from panels **A,D**. As depicted in these length history schematics, microtubules and some other polymers (e.g., PhuZ, ParM) display a behavior called dynamic instability (DI), in which individual filaments alternate stochastically between periods of growth and shortening (Mitchison & Kirschner, 1984; Erb et al., 2014; Garner, Campbell, & Mullins, 2004). Transitions from growth to shortening are called catastrophes (label *C* in panels **C,F**). Transitions from shortening to growth are called rescues (label *R* in panels **C,F**), *if* the filament has not completely depolymerized (label *D* in panels **C,F**). ***Significance for filament and population behaviors***: When *J* > 0 in a dilution or constant [free subunit] experiment, the average filament length of the population increases over time (reaches a polymer-growth steady state where the rate of increase is constant with time), and individual filaments experience net growth over time. When *J* < 0 in a dilution experiment, the average filament length decreases over time, and individual filaments experience net shortening over time. When *J* = 0 in a constant [free subunit] experiment, the average filament length increases initially and then reaches a plateau over time (this is polymer-mass steady state); individual filaments repeatedly grow and completely depolymerize, but experience no net change in length over sufficiently long time periods. Note that panels **E-F** are analogous to Figure 1 of (Dogterom & Leibler, 1993).

When the net flux of subunits into a population’s polymer mass is *positive* (*J* > 0), the average filament length *increases* over time (Figure 1B,E, squares). In contrast, when the net flux of subunits into a population’s polymer mass is *negative* (*J* < 0), the average filament length *decreases* over time (Figure 1B, circles). Polymer-mass steady state is when the net flux of subunits into a population’s polymer mass is *zero* (*J* = 0); in this situation, the average filament length stays *constant* over time (Figure 1E, diamonds). The [free subunit] above which *J* > 0 is a critical concentration (CC). This CC can be described as the [free subunit] above which “net assembly” (Walker et al., 1988) or “unbounded growth” (Dogterom & Leibler, 1993; Fygenson, Braun, & Libchaber, 1994) will occur (CC_NetAssembly_ in Figure 2) (see also (Hill & Chen, 1984; Hill, 1987; Jonasson et al., 2019).

**Figure 2:**
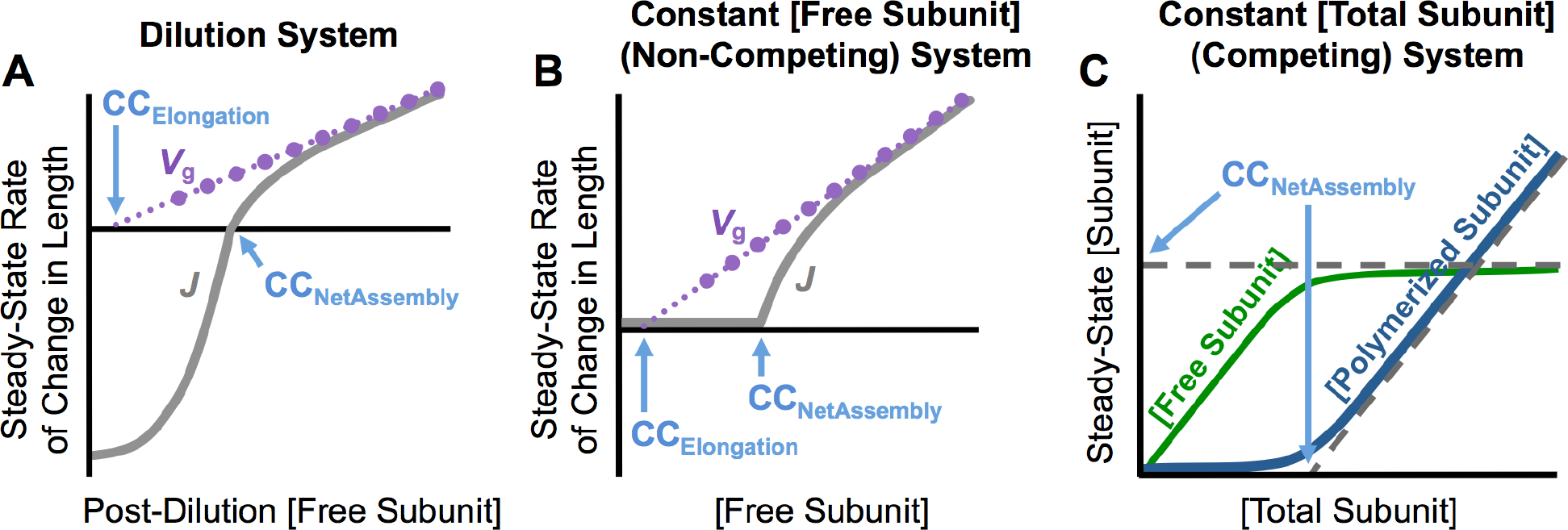
Schematic representation of critical concentrations. **(*A*)** Overlay of *V*_g_ and *J* as functions of post-dilution [free subunit] in a dilution experiment. **(*B*)** Overlay of *V*_g_ and *J* as functions of [free subunit] in a constant [free subunit] experiment. *V*_g_ is the growth velocity of individual filaments during the growth phase of dynamic instability. *J* is the flux of subunits into and out of polymer *or* the rate of change in average filament length of a population; *J* encompasses both growth and shortening phases. **(C)** Plots of steady-state [free subunit] and [polymerized subunit] as functions of [total subunit] in a competing system. ***Definitions of critical concentrations (CCs)***: The [free subunit] above which *V*_g_ > 0 is CC_Elongation_. This CC is determined by plotting *V*_g_ as a function of [free subunit] and extrapolating back to the [free subunit] at which *V*_g_ = 0 (panels **A-B**). The [free subunit] above which *J* > 0 is CC_NetAssembly_. This CC can be determined from plotting *J* as a function of [free subunit] and identifying the [free subunit] at which *J* crosses zero in a dilution experiment (panel **A**) or at which *J* transitions from zero to positive in a constant [free subunit] experiment (panel **B**). CC_NetAssembly_ as measured from *J* (in either panel **A** or **B**) is equivalent to the CC as measured in a competing system (panel **C**).

### 1.2 Dynamic instability of individual microtubules

Dynamic instability (DI) is a behavior in which individual microtubules stochastically alternate between periods of growth and shortening (Mitchison & Kirschner, 1984) (Figure 1C,F). Transitions from growth to shortening are called catastrophes. Transitions from shortening to growth, without complete depolymerization, are called rescues. DI is commonly quantified by four parameters: growth velocity (*V*_g_), shortening velocity (*V*_s_), catastrophe frequency (*F*_cat_), and rescue frequency (*F*_res_). The [free tubulin] above which *V*_g_ > 0 is the CC above which “elongation” phases of individual filaments can occur (CC_Elongation_ in Figure 2A-B) (Hill & Chen, 1984; Hill, 1987; Walker et al., 1988; see also Jonasson et al., 2019).

### 1.3 Relationship between flux and dynamic instability

Individual MTs can display DI when *J* is positive, negative, or zero. When *J* is *positive*, individual MTs experience *net growth* (more length increase during growth than length decrease during shortening) over sufficient time (Figure 1C,F, label *J* > 0). When *J* is *negative*, individual MTs experience *net shortening* (more length decrease during shortening than length increase during growth) over sufficient time (Figure 1C, label *J* < 0). When *J* equals *zero*, individual MTs experience *no net length change* over sufficient time (Figure 1F, label *J* = 0); in this case, growth and shortening can both occur, but the length changes balance each other out.

### 1.4 Equation relating flux and dynamic instability

Previous papers have presented various forms of an equation relating dynamic instability to the flux of subunits into and out of polymer. To our knowledge, this equation was first presented in (Hill & Chen, 1984). The flux of subunits into and out of polymer for an individual microtubule over sufficient time or averaged over MTs in a sufficiently large population can be given by

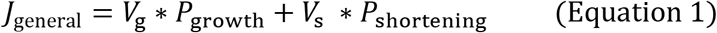

(Hill & Chen, 1984; Komarova et al., 2002). *V*_g_ is the growth velocity during growth phases and *V*_s_ is the shortening velocity during shortening phases. *P*_growth_ and *P*_shortening_ are the probabilities of being in growth or shortening. *P*_growth_ can be thought of as the proportion of time in growth (Hill & Chen, 1984; Walker et al., 1988; Gliksman, Parsons, & Salmon, 1992) or the proportion of individuals that are in growth within a population (Komarova et al., 2002), and analogously for *P*_shortening_. Note that *J* can be determined from *V*_g_, *V*_s_, *P*_growth_, and *P*_shortening_ by using Equation 1, but that *V*_g_, *V*_s_, *P*_growth_, and *P*_shortening_ cannot be uniquely determined from *J* alone.

We use *V*_s_ to mean the shortening *velocity* (negative number). If *V*_s_ is used to mean the shortening *speed* (positive number), then the plus sign in Equation 1 would be become a minus sign: *J* = *V*_g_ ∗ *P*_growth_ − *V*_s_ ∗ *P*_shortening_. These sign conventions are chosen so that growth results in an increase in length and shortening results in a decrease in length.

Under conditions where *P*_shortening_ is near zero (e.g., if [free subunit] is high), *J* is approximately equal to *V*_g_ (Figure 3, dotted line). In this case, almost all individual MTs in a population would be in growth. Under conditions where *P*_growth_ is near zero (e.g., if [free subunit] is low), *J* is approximately equal to *V*_s_ (Figure 3, dashed line). In this case, almost all individual MTs in a population would be in shortening.

**Figure 3:**
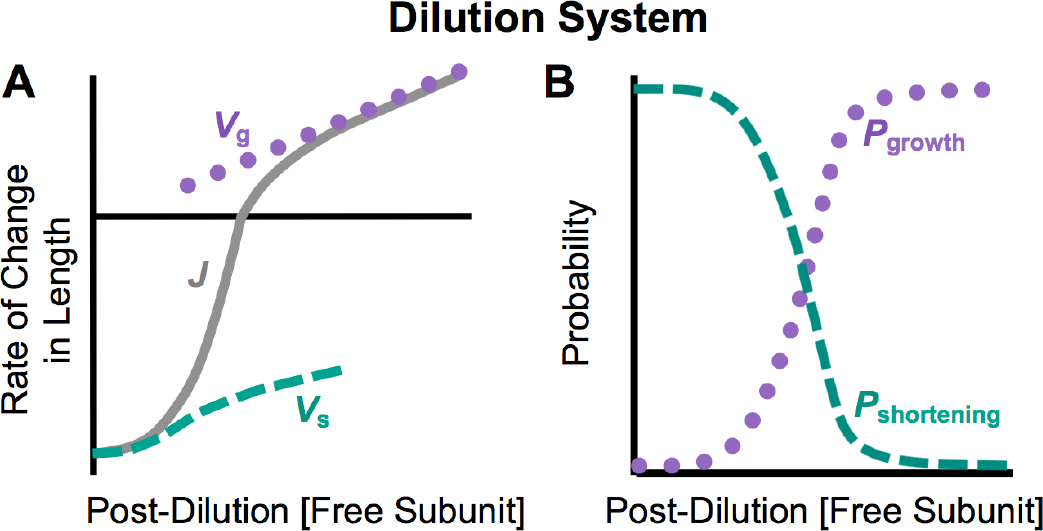
Schematic representation of net flux and the velocities and probabilities of growth and shortening in a dilution experiment. This figure summarizes predictions of existing models in the literature (see Figure 8 of (Bayley, Schilstra, & Martin, 1990) and Figure 4 of (Hill & Chen, 1984)). **(*A*)** *J* (net flux), *V*_g_ (growth velocity), and *V*_s_ (shortening velocity) as functions of post-dilution [free subunit]. **(*B*)** *P*_growth_ (probability of growth) and *P*_shortening_ (probability of shortening) as functions of post-dilution [free subunit]. ***Significance***: The *J*_General_ equation (Equation 1) relates *J* to *V*_g_, *V*_s_, *P*_growth_, and *P*_shortening_: *J*_General_ = *V*_g_ ∗ *P*_growth_ + *V*_s_ ∗ *P*_shortening_. At sufficiently high [free subunit], all or almost all individual filaments are growing (*P*_growth_ ≈ 1, panel **B**), so the net flux and the growth velocity are approximately equal (*J* ≈ *V*_g_, panel **A**). Reciprocally, at sufficiently low [free subunit], all or almost all individual filaments are shortening (*P*_shortening_ ≈ 1, panel **B**), so the net flux and the shortening velocity are approximately equal (*J* ≈ *V*_s_, panel **A**). For intermediate concentrations, growth and shortening coexist (*P*_growth_ and *P*_shortening_ > 0, panel **B**). As [free subunit] increases, *J* increases from *V*_s_ to *V*_g_ (panel **A**) as *P*_shortening_ decreases from 1 to 0 and *P*_growth_ increases 0 to 1 (panel **B**). This cartoon depicts an increasing and non-linear dependence of *V*_s_ on [free subunit], as predicted by previous models (see Figure 8 of (Bayley, Schilstra, & Martin, 1990) and Figure 4 of (Hill & Chen, 1984).

For a population of MTs, *J* as written in Equation 1 represents the per microtubule average flux of subunits into and out of polymer. For all equations in this paper we assume that *V*_g_, *V*_s_, and *J* are in units of length/time (e.g., µm/s). In this case, *J* is equivalent to the rate of change in average MT length (in this paper, our average MT length calculated as the sum of the lengths of all individuals MTs in the population divided by the number of stable MT seeds). If the right hand side of Equation 1 (or any of the subsequent *J* equations in this paper) is multiplied by the number of individuals in a population and the units are converted to concentration/time (e.g., µM/s), then *J* will represent the flux of subunits into and out of the population’s overall polymer mass, instead of the per microtubule average length. The quantity *J* in Equation 1 is also referred to as the drift coefficient, which represents the rate of displacement of the MT ends (Vorobjev, Rodionov, Maly, & Borisy, 1999; Maly, 2002; Komarova et al., 2002; Vorobjev & Maly, 2008; Mirny & Needleman, 2010).

For a system where both ends (plus and minus) of each filament are free, Equation 1 can be applied to each end separately. In this case, the rate of change in average MT length would be the sum of *J* for the plus end and *J* for the minus end. If one end of each filament is anchored (e.g., at a centrosome), then *J* for the free end is equivalent to the rate of change in average MT length. The work in this paper examines the latter case, in which MTs are active at only one end.

### 1.5 Role of flux and dynamic instability in defining critical concentrations

The relationship between *J* and *V*_g_ in the *J* equation can be useful in understanding two critical concentrations (CC_NetAssembly_ and CC_Elongation_ in Figure 2) that are relevant to the behaviors of DI polymers. CC_NetAssembly_ (called *c*_o_ in (Hill & Chen, 1984), *c*_cr_ in (Dogterom & Leibler, 1993), CC_PopGrow_ in (Jonasson et al., 2019)) is the higher of these two CCs and is the [free tubulin] above which *J* > 0. At [free tubulin] above CC_NetAssembly_, the average MT length or polymer mass of a population will increase persistently, and individuals will experience net growth over time.

CC_Elongation_ (called *c*_1_ in (Hill & Chen, 1984), *S*_c_^e^ in (Walker et al., 1988), CC_IndGrow_ in (Jonasson et al., 2019)) is the [free tubulin] above which *V*_g_ > 0. CC_Elongation_ is measured by extrapolation to *V*_g_ = 0 from a plot of *V*_g_ versus [free tubulin]. At [free tubulin] above CC_Elongation_, individual MTs can exhibit transient growth phases, though for [free tubulin] near CC_Elongation_, few MTs will exceed experimentally relevant detection thresholds (Jonasson et al., 2019).

For polymers that do not display (detectable) DI, CC_Elongation_ and CC_NetAssembly_ are either the same (e.g., equilibrium polymers) or experimentally indistinguishable (e.g., actin). For such polymers, when *J* > 0, individuals grow (*P*_growth_ ≈ 1) and *J* ≈ *V*_g_; when J < 0, individuals shorten (*P*_shortening_ ≈ 1) and *J* ≈ *V*_s_. In contrast, for polymers that display DI, CC_Elongation_ and CC_NetAssembly_ are distinguishable (Jonasson et al., 2019). This is case depicted in the schematics in Figures 1 to 3, and that will be examined in this paper.

### 1.6 Outline of approach

In this paper, we compare variants of the *J* equation, examine conditions under which the different forms of the equation are equivalent, and demonstrate how to convert between the forms. We will use subscripts on *J* to distinguish the specific versions of the equation. As the starting point for the analysis, we use the *J*_General_ equation (Equation 1), which depends on the probabilities of being in growth or shortening, *P*_growth_ and *P*_shortening_, respectively (similar to (Hill & Chen, 1984; Komarova et al., 2002)). In the Results and Discussion section, we first examine forms of the equation (*J*_Time_ and *J*_TimeStep_) in which *P*_growth_ and *P*_shortening_ are determined from the fraction of time spent in growth or shortening (e.g., (Komarova et al., 2002)). Next, versions of the equation (*J*_DI_ and *J*_DI_piecewise_) that use *F*_cat_ and *F*_res_ to calculate *P*_growth_ and *P*_shortening_ are considered (Hill & Chen, 1984; Walker et al., 1988; Verde et al., 1992; Dogterom & Leibler, 1993).

The *J*_DI_ form of the equation is perhaps the most well-known, because Dogterom and colleagues related this form of the equation to bounded and unbounded growth behaviors (Verde et al., 1992; Dogterom & Leibler, 1993). In the “bounded” growth regime, the average MT length reaches a steady-state value over time. In the “unbounded” growth regime, the average MT length increases indefinitely. Forms of the *J* equation, most commonly *J*_DI_, have been utilized in many other papers (e.g., (Gliksman et al., 1992; Bicout, 1997; Maly, 2002; Vorobjev & Maly, 2008; Mirny & Needleman, 2010; Yarahmadian, Barker, Zumbrun, & Shaw, 2011; Mahrooghy, Yarahmadian, Menon, Rezania, & Tuszynski, 2015; Ishihara et al., 2016; Aparna, Padinhateeri, & Das, 2017; Lamson, Edelmaier, Glaser, & Betterton, 2019; Kuo, Trottier, Mahamdeh, & Howard, 2019)).

We use our previously established computational simulations to illustrate the results of the *J* equations at various tubulin concentrations and to demonstrate how aspects of experimental design, such as the timing of the experimental steps, can lead to errors in measuring *J*.

As discussed above, the variants of the *J* equation use measurements on individuals to calculate estimates of *J*. To assess these variants, we compared each to *J*_Net_, which is the *net* rate of change in average MT length as calculated directly from the change in the average MT length of the population between two time points. *J*_Net_ provides the true net rate of change that occurs in any particular run of the simulation, and is therefore a useful baseline for comparison.

## 2 RESULTS AND DISCUSSION

### 2.1 Computational Simulations

In our dimer-scale computational model (introduced in (Margolin, Goodson, & Alber, 2011; Margolin et al., 2012)), the attachment and detachment of subunits from protofilaments, the formation and breakage of lateral bonds between protofilaments, and the hydrolysis of subunits from GTP-tubulin to GDP-tubulin are modeled as discrete events. The biochemical kinetic rate constants for these processes are inputted by the user. The values of the rate constants used here were tuned in (Margolin et al., 2012) to approximate the plus-end dynamics of mammalian MTs as reported in (Walker et al., 1988). The MTs grow from stable non-hydrolyzable GTP-tubulin seeds, and all attachment and detachment events occur at the free end of each MT. The values of *J* and the DI parameters are emergent properties of the system. This is analogous to experimental systems, where the values of *J* and the DI parameters will depend on cell type and experimental conditions such as the source of the tubulin and the buffer. For additional details about the simulations, please see the Methods, Section 3.1.

We use our computational model to simulate dilution experiments and constant [free tubulin] experiments (see Figure 1 for schematic representations of *J* and DI behaviors in these types of experiments). Except where otherwise indicated, all simulations in this paper were performed with a population of 50 stable MT seeds.

### 2.2 Time-based *J* Equation

One way to calculate *P*_growth_ and *P*_shortening_ to use in Equation 1 is

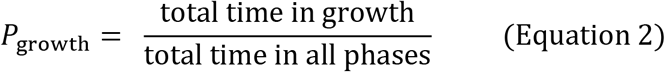

and

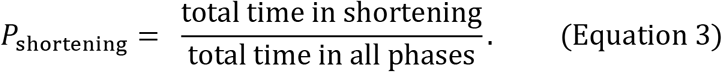

Then *J* can be calculated as

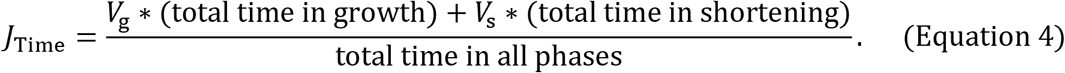

One method for obtaining values of *V*_g_, *V*_*s*_, total time in growth, and total time in shortening is to use standard DI analysis to identify periods of growth and shortening in length histories (see Methods, Section 3.2.2). Figure 4 shows plots of *V*_g_, *V*_*s*_ (panels **A,C**), *P*_growth_, and *P*_shortening_ (panels **B,D**) as obtained from the DI analysis, as well as *J*_Net_ for comparison (panels **A,C**). At very high [free tubulin] in both the dilution and constant [free tubulin] simulations, *P*_shortening_ ≈ 0, *P*_growth_ ≈ 1, and J ≈ *V*_g_. In contrast, in the dilution simulations, at very low post-dilution [free tubulin], *P*_shortening_ ≈ 1, *P*_growth_ ≈ 0, and *J* ≈ *V*_s_. However, in the constant [free tubulin] simulations, at very low [free tubulin], *P*_shortening_ ≈ *P*_growth_ ≈ 0, and J ≈ 0.

**Figure 4:**
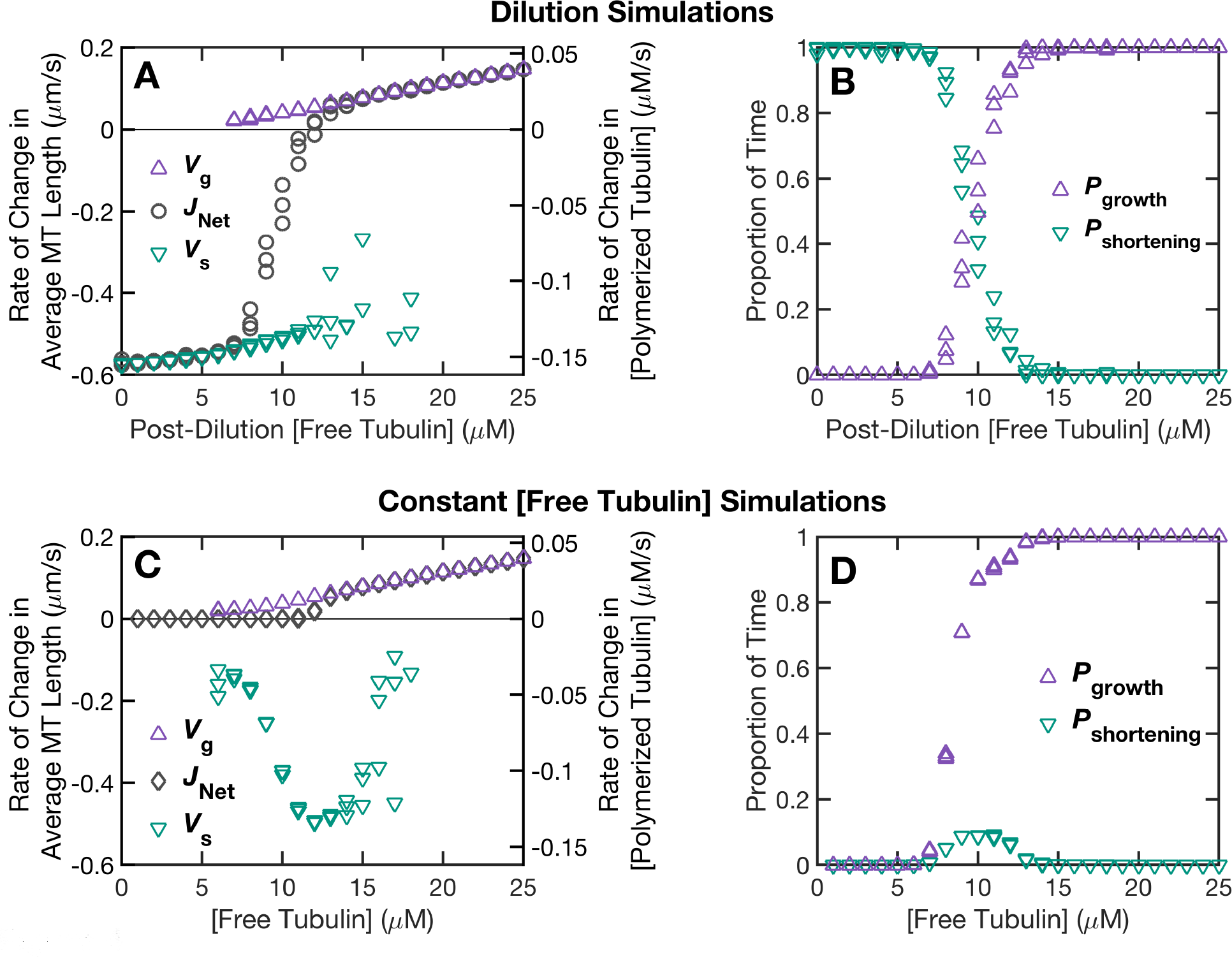
Relationship between net flux and the velocities and probabilities of growth and shortening as observed in the simulation data. Top row: dilution simulations; bottom row: constant [free tubulin] simulations. **(*A,C*)** Overlay of *J*_Net_, *V*_g_, and *V*_s_ as functions of [free tubulin]. **(*B,D*)** *P*_growth_ (proportion of time in growth) and *P*_shortening_ (proportion of time in shortening) as functions of [free tubulin]. ***Methods***: In the dilution simulations, the pre-dilution [total tubulin] was 20 µM, and the dilution into new [free tubulin] was performed at minute 5 of the simulation (panels **A-B**). *J*_Net_ is the net rate of change in average MT length of the population (Equation M1 of the Methods, Section 3.2.1):

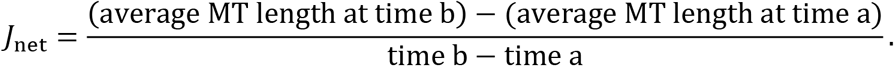 In panels **A-B**, time a = 5 seconds after the dilution and time b = 40 seconds after the dilution. In panels **C-D**, time a = minute 15 and time b = minute 30 of the simulations. The effect of varying the measurement times will be examined in Sections 2.6 and 2.7. The measurements of *V*_g_, *V*_s_, and the times in growth and shortening were obtained using the DI analysis method described in the Methods, Section 3.2.2; these measurements were taken during the same time periods used to obtain *J*_Net_. In the DI analysis, we imposed a threshold of 25 subunit lengths (200 nm) on the length change that must occur for a growth or shortening segment to be detected. This threshold was chosen to be comparable to typical experimental detection limits in fluorescence microscopy. Data points are plotted for each of three independent replicates of the simulations at each value of (post-dilution or constant) [free tubulin]. ***Interpretations***: Our dilution simulation results (panels **A-B**) are consistent with the predictions of existing models for dilution experiments (Figure 3). The results in panels **C-D** are from simulations of a different type of experiment, specifically, a constant [free tubulin] experiment. The differences between dilution experiments and constant [free tubulin] experiments occur because the MTs in dilution experiments are very long at the time of the dilution (allowing *J* to be negative after the dilution), whereas MTs in constant [free tubulin] experiments can only depolymerize as far as they have grown (*J* cannot be negative). In the [free tubulin] range where *J* > 0 (i.e., above CC_NetAssembly_, Figure 2), the results from the dilution simulations and the constant [free tubulin] simulations are similar to each other. In contrast, in the [free tubulin] range below CC_NetAssembly_, *J* < 0 in the dilution simulations (panel **A**) and *J* = 0 in the constant [free tubulin] simulations (panel **C**).

The difference at low [free tubulin] between the dilution and constant [free tubulin] systems occurs because the MTs in dilution systems are sufficiently long as to not undergo complete depolymerizations back to the seed during the measurement period (Figure 5A-B); in contrast, the MTs in constant [free tubulin] systems are short at low tubulin and therefore frequently and repeatedly depolymerize back to the seed (Figure 5C-D).

**Figure 5:**
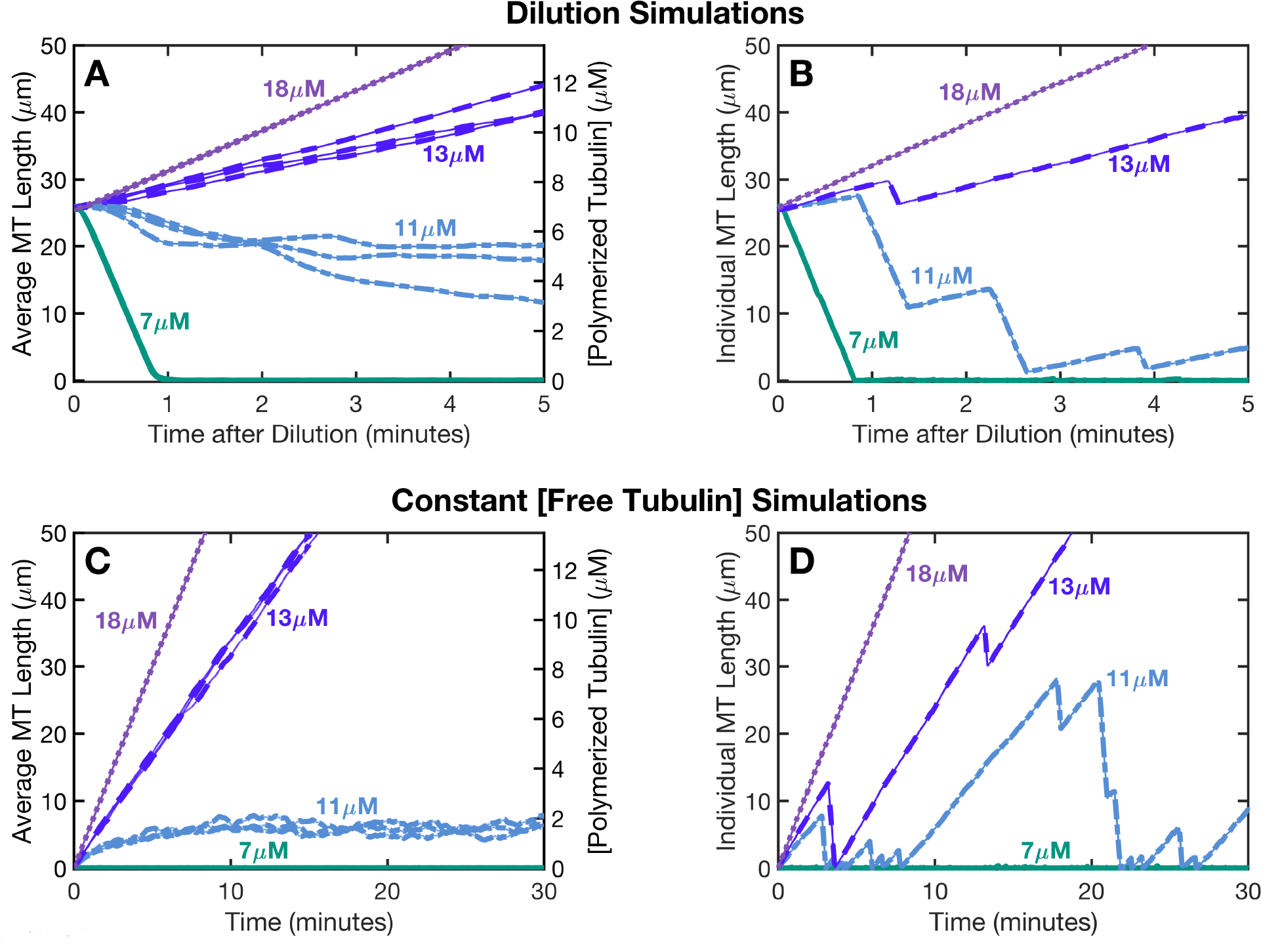
Average and individual MT lengths versus time. Top row: dilution simulations; bottom row: constant [free tubulin] simulations. **(*A,C*)** Average length of the MT population (left axis) or [polymerized tubulin] (right axis) versus time for the [free tubulin] concentrations as indicated on the plots. At each [free tubulin] shown here, the average MT length is plotted for three independent replicates (same simulation runs used in Figure 4). **(*B,D*)** Individual MT length versus time for a representative individual from each [free tubulin] in panels **A,C**. ***Methods***: In the dilution simulations (panels **A-B**), the pre-dilution [total tubulin] was 20 µM and the dilution into new [free tubulin] was performed at minute 5 of the simulation. Under these conditions, the lengths of almost all individual MTs in the population at the time of the dilution were between ~25 and ~27 µm. The dilution simulations were run for 5 minutes after the dilution time. The constant [free tubulin] simulations (panels **C-D**) were run for 30 minutes. ***Interpretations:** Behaviors of populations (left column) and individuals (right column):* In both the dilution and constant [free tubulin] simulations, when [free tubulin] is above ~12 µM (CC_NetAssembly_), the average MT length increases over time (*J* > 0) (panels **A,C**), and individual MTs experience net growth over time (panels **B,D**). In this [free tubulin] range, the system reaches a polymer-growth steady state where *J* is constant over time. Conversely, below ~12 µM, the dilution and constant [free tubulin] simulations differ from each other. In the dilution simulations below ~12 µM, the average MT length decreases over time (*J* < 0) (panel **A**), and individual MTs experience net shortening over time (panel **B**). In the constant [free tubulin] simulations below ~12 µM, the average MT length levels off over time (*J* = 0, polymer-mass steady state) (panel **C**), and individual MTs experience no net change in length over sufficient time (panel **D**). Note that growth and shortening phases of individual MTs can occur for *J* > 0, *J* = 0, and *J* < 0. *Relationship to J measurement periods*: For the dilution simulations (panels **A-B**), the measurements of *J* in Figure 4A were taken from 5 to 40 seconds after the time point of the dilution. Based on the MT length data (panels **A-B**), this measurement period was chosen to avoid complete MT depolymerizations. For example, as seen in panel **A** for 7 µM, the average MT length decreased to ~0 µm by ~1 minute after the dilution, meaning that all individuals in the population had completely depolymerized. For the constant [free tubulin] simulations (panels **C-D**), the measurements of *J* in Figure 4C were taken from minute 15 to minute 30 of the simulations, chosen to be after the system had reached the appropriate steady state (either polymer-growth or polymer-mass steady state, depending on the [free tubulin]). For example, in panel **C**, the average MT length for 11 µM levels off after ~10 minutes.

Figure 6 shows the results of the *J*_Time_ equation (Equation 4) evaluated with *V*_g_, *V*_*s*_, total time in growth, and total time in shortening as measured with our DI analysis method (Methods, Section 3.2.2). The *J*_Time_ results match well with the direct measurements of *J* from the net rate of change in average MT length, *J*_Net_, in the simulations data (Figure 6).

**Figure 6:**
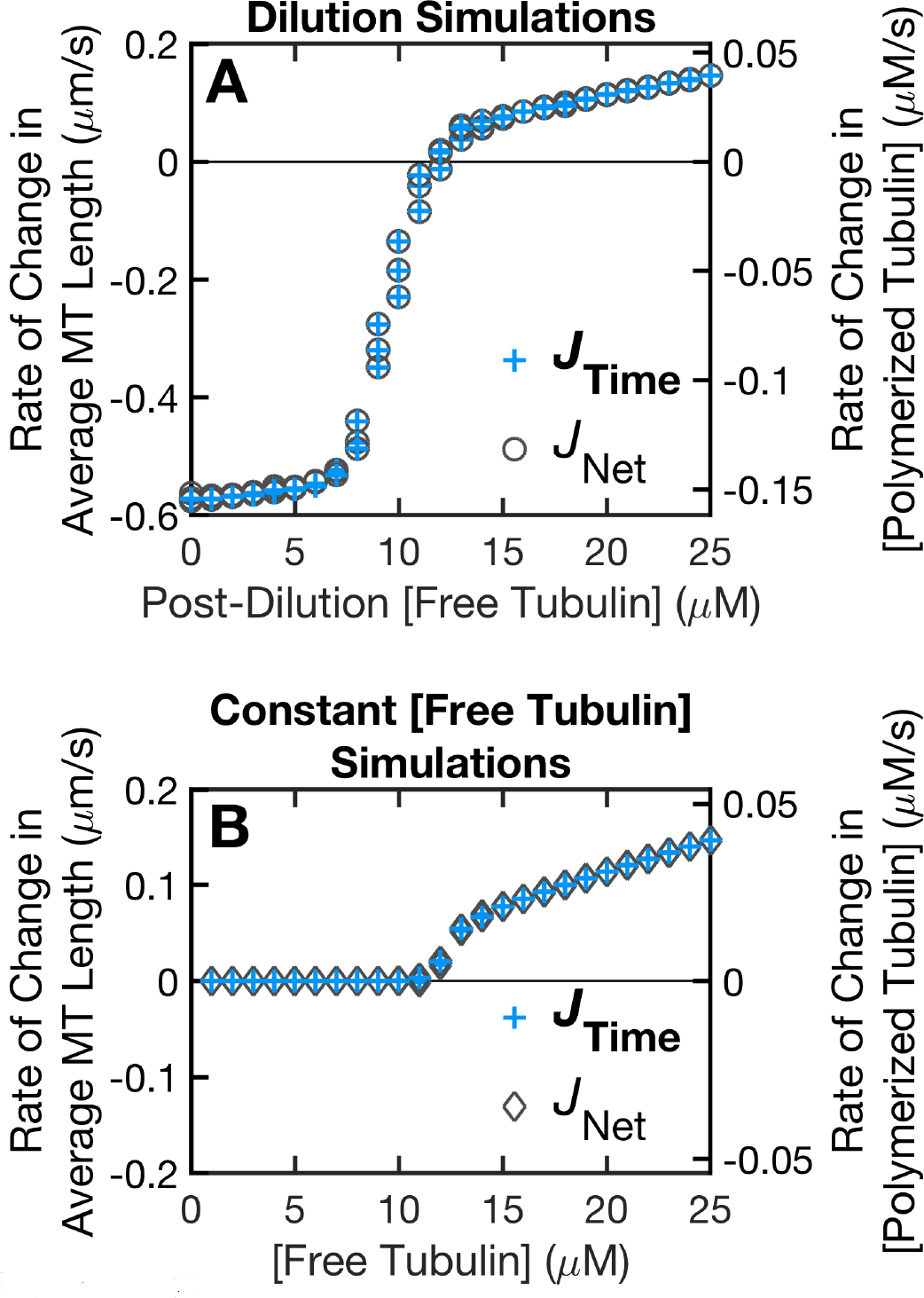
Comparison of the *J*_time_ equation to the *J*_Net_ data. For **(*A*)** the dilution simulations and **(*B*)** the constant [free tubulin] simulations, this figure shows a comparison of the results of the *J*_Time_ equation (+ symbols; Equation 4) to *J*_Net_ (circle symbols in panel **A**, diamond symbols in panel **B**), which is the net rate of change in average MT length (left axes) or in polymer mass (right axes). ***Methods***: The *J*_Net_ data are re-plotted from Figure 4. The *J*_Time_ equation,

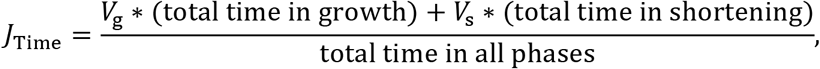

is evaluated with measurements of *V*_g_, *V*_s_, and the total times obtained using the DI analysis method described in the Methods, Section 3.2.2 (same DI measurements as in Figure 4). The DI analysis calculates *V*_g_ as (total length change during growth phases) / (total time in growth) and *V*_s_ as (total length change during shortening phases) / (total time in shortening). Data points are plotted for each of three independent replicates of the simulations at each value of (post-dilution or constant) [free tubulin]. ***Interpretations***: The data show that the *J*_Time_ equation evaluated with these DI measurements matches well with the *J*_Net_ data over the full range of concentrations. If *V*_g_ and *V*_s_ were calculated by a different method (e.g., fitting a regression line to each growth or shortening segment, and then averaging the slopes over all growth segments or all shortening segments), then the results of the *J*_Time_ equation could deviate slightly from the net rate of change data.

When applying the *J*_Time_ equation (Equation 4), one should be aware of the possibility of other states besides growth and shortening. In Equations 2, 3, and 4, “total time in all phases” is the time of all observations and can include time in other phases (e.g., pause) during which the MT length is approximately constant. Thus, total time may be greater than (total time in growth) + (total time in shortening). If (total time in growth) + (total time in shortening) were used in place of total time in Equation 4 and if the MTs spent some time in an additional state such as pause, then the equation would overestimate the magnitude of the actual rate of change in average length. However, the sign of *J* would not be affected, so the equation could still be used to determine if the average length was increasing (*J* > 0), decreasing (*J* < 0), or constant (*J* = 0).

### 2.3 Time-step method for measuring *J*

An alternative approach to determine the value of *J* (called the drift coefficient and abbreviated as *v*_d_ in (Komarova et al., 2002)) is to measure the displacements of the MT ends over short time steps (e.g., between successive images in a time series); then

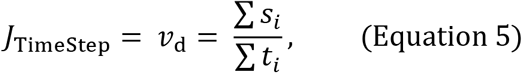

where **∑***s*_*i*_ is the sum of all the displacements of MT ends and **∑***t*_*i*_ is the sum of the corresponding time changes (Komarova et al., 2002). Figure 7 compares the results of the *J*_TimeStep_ equation and the *J*_Net_ data (*J*_Net_, as above, is measured directly from the net rate of change in average MT length); the results of the two methods agree well. For implementation details of the time-step analysis method, please see the Methods, Section 3.2.3.

#### 2.3.1 Time steps during which a displacement is zero (*s*_*i*_ = 0)

A factor to be aware of when measuring *J* by the *J*_TimeStep_ approach is whether the experimental method can track displacements of zero (*s*_*i*_ = 0). To analyze this situation, let **∑**_pos_, **∑**_neg_, and **∑**_zero_, respectively, represent sums (of displacements, or of the corresponding times) when the displacements are positive (*s*_*i*_ > 0), negative (*s*_*i*_ < 0), or zero (*s*_*i*_ = 0). For example, **∑**_zero_ *t*_*i*_ represents the sums of time steps during which the displacement is zero (*s*_*i*_ = 0). Then **∑**_zero_ *s*_*i*_ = 0, so **∑***Si* = **∑**_pos_ *s*_*i*_ + **∑**_neg_ *s*_*i*_; in other words, any displacements equaling zero (*s*_*i*_ = 0) do not affect the value of **∑***s*_*i*_. However, **∑***t*_*i*_ = **∑**_pos_ *t*_*i*_ + **∑**_neg_ *t*_*i*_ + **∑**_zero_ *t*_*i*_, so **∑**_zero_ *t*_*i*_ can affect **∑***t*_*i*_. If the experimental method used does not detect displacements of zero, then **∑***t*_*i*_ may be underestimated and therefore the magnitude of *J* would be overestimated. The relevance to any specific system would depend on whether there are displacements of zero and how often they occur.

**Figure 7:**
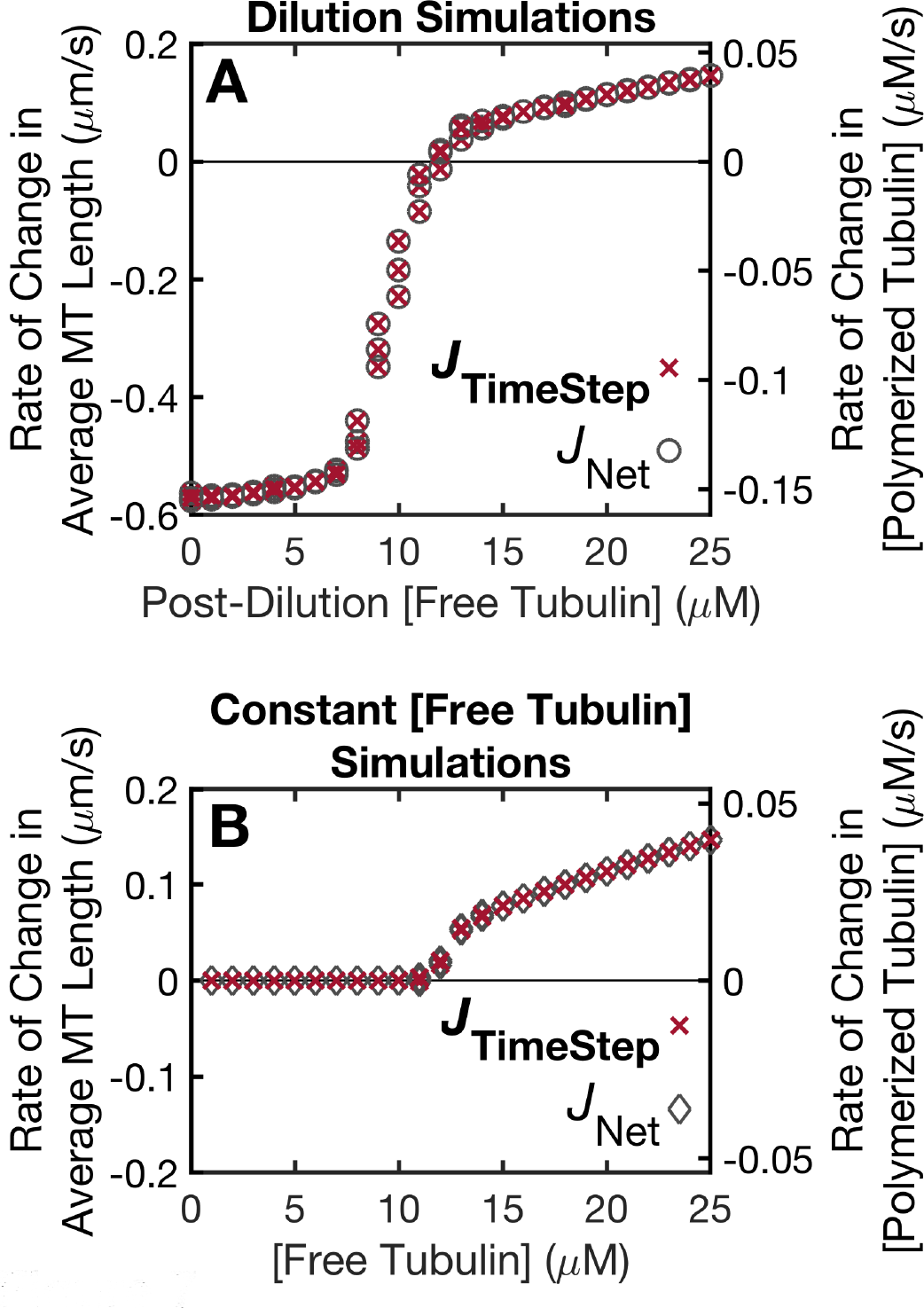
Comparison of the *J*_TimeStep_ equation to the *J*_Net_ data. For **(*A*)** the dilution simulations and **(*B*)** the constant [free tubulin] simulations, this figure shows a comparison of the results of the *J*_TimeStep_ equation (x symbols; Equation 5) to *J*_Net_ (circle symbols in panel **A**, diamond symbols in panel **B**). ***Methods***: The *J*_Net_ data are re-plotted from Figure 4. To evaluate the *J*_TimeStep_ equation (called *v*_d_ or drift coefficient in (Komarova et al., 2002)),

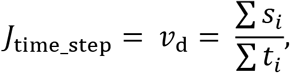

the length history of each MT is divided into short time steps (*t*_i_), here 1 second, and the displacement (*s*_i_) of the MT end over each time step is recorded. These measurements were performed during the same time period as the *J*_Net_ measurements (minute 15 to minute 30 in the constant [free tubulin] simulations; 5 to 40 seconds after the time of dilution in the dilution simulations). The displacements and corresponding time steps were then summed over all individuals and over the total measurement period. For additional information about this time-step analysis method, please see the Methods, Section 3.2.3. Data points are plotted for each of the three independent replicates of the simulations at each value of (post-dilution or constant) [free tubulin]. ***Interpretations***: The data show that the *J*_TimeStep_ equation evaluated with the displacement measurements matches well with the *J*_Net_ data over the full range of concentrations.

#### 2.3.2 Mathematical equivalence of *J*_TimeStep_ (Equation 5) to *J*_General_ (Equation 1) and *J*_Time_ (Equation 4)

To see the equivalence of *J*_Time_ (Equation 4) and *J*_TimeStep_ (Equation 5), the displacements can be separated into positive and negative displacements. *V*_g_ and time in growth are determined from the positive displacements, while *V*_s_ and time in shortening are determined from the negative displacements. More specifically, *V*_g_ is **∑**_pos_ *s*_*i*_ / **∑**_pos_ *t*_*i*_ where the sums include only the positive displacements;**∑**_pos_ *t*_*i*_ is the time in growth; then

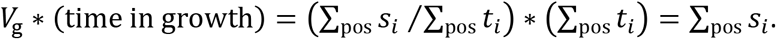

Similarly, *V*_s_ = **∑**_neg_ *s*_*i*_ / **∑**_neg_ *t*_*i*_ where the sums include only the negtive displacements; **∑**_neg_ *t*_*i*_ = time in shortening; and

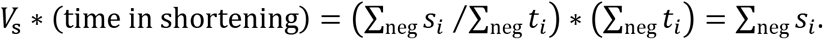

Also, **∑** *s*_*i*_ = **∑**_pos_ *s*_*i*_ + **∑**_neg_ *s*_*i*_. Then,

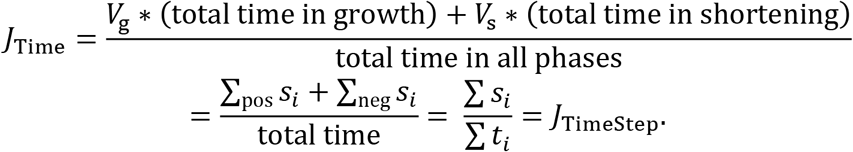

Thus, *J*_TimeStep_ (Equation 5) is algebraically equivalent to *J*_Time_ (Equation 4), and therefore to *J*_General_ (Equation 1).

Alternatively, the equivalence of *J*_General_ (Equation 1) and *J*_TimeStep_ (Equation 5) can be shown using *P*_growth_ = **∑**_pos_ *t*_*i*_ / **∑** *t*_*i*_ and *P*_shortening_ = **∑**_neg_ *t*_*i*_ / **∑***t*_*i*_, yielding

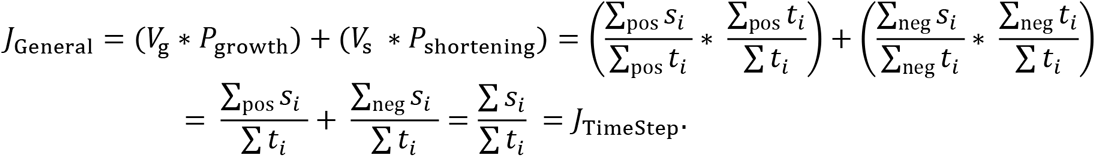

Note that *V*_g_ and *V*_s_ as calculated from the time-step method depend on the size of the time-step (see Supplement Methods of (Jonasson et al., 2019)), and may differ from *V*_g_ and *V*_s_ as calculated from the DI analysis method described above (Section 2.2). However, the results of the *J*_Time_ or *J*_TimeStep_ equations will still fit well with the data, as long as the velocities and the probabilities are determined in a way that is internally consistent (Figure 8).

**Figure 8:**
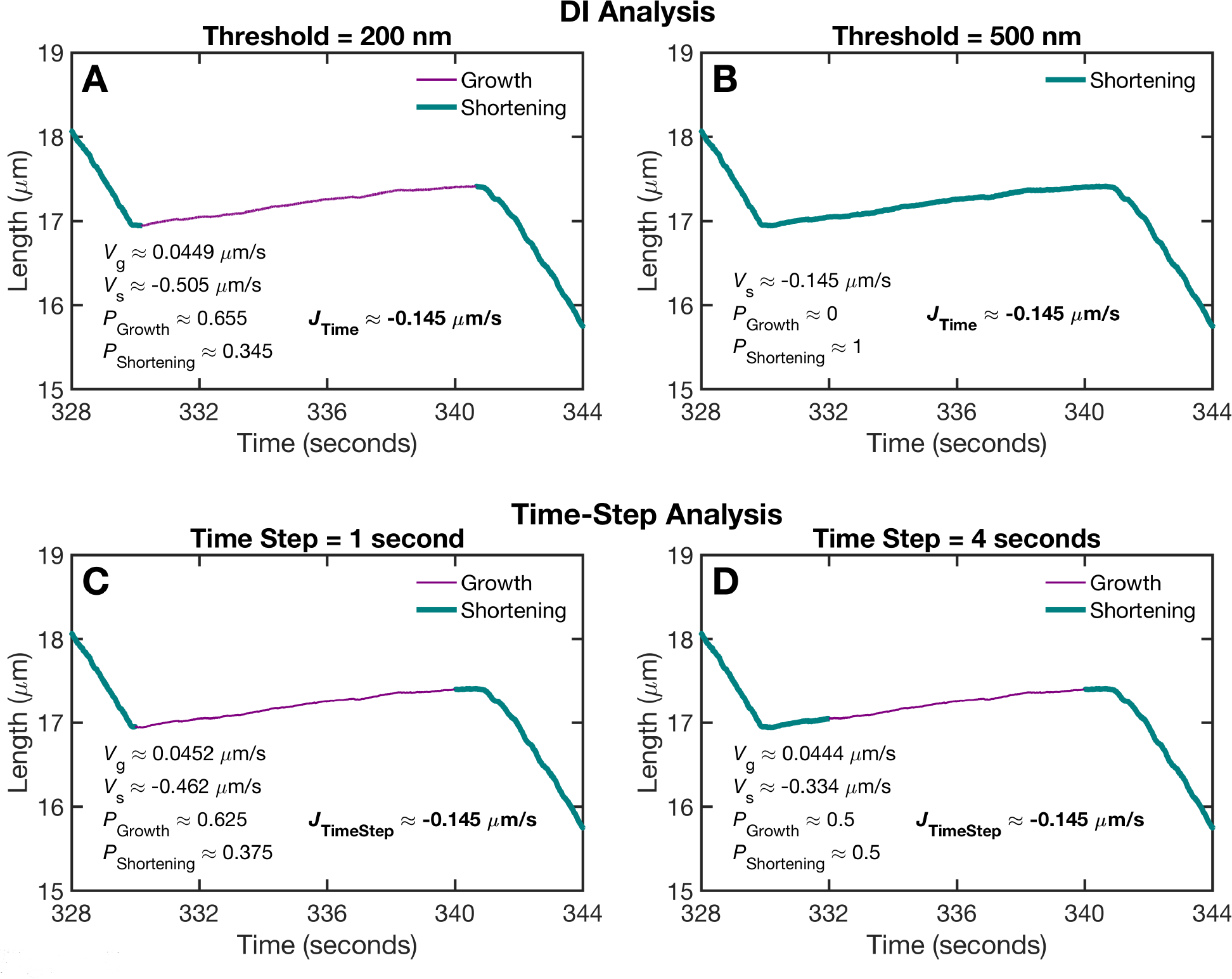
Comparison of measurements obtained from the DI analysis and the time-step analysis methods. **(*A,B*)** Dynamic instability analysis (as described in the Methods, Section 3.2.2) with a threshold of either 200 nm (panel **A**) or 500 nm (panel **B**) for the length change that must occur for a growth or shortening segment to be detected. **(*C,D*)** Time-step analysis (as described in the Methods, Section 3.2.3) with a time-step of either 1 second (panel **C**) or 4 seconds (panel **D**). ***Methods***: The panels show a portion of a particular length history from the dilution simulations at 11 µM, as an example to illustrate the analysis methods. The numerical values on each plot are based only on the portion of the length history shown. Note that the analysis results in the other figures are based on all individuals in a population and use longer time periods. For each of the two methods, *V*_g_, *V*_s_, *P*_growth_ and *P*_shortening_ were calculated as described in the main text (Sections 2.2, 2.3, and 3.2.2). ***Interpretations***: How a length history is divided into growth and shortening segments differs between the two analysis methods (compare top and bottom rows). Additionally, within either method, the results differ based on the size of the threshold (compare panels **A** and **B**) or the size of the time step (compare panels **C** and **D**). Since the division of the length history into growth and shortening differs among the panels, the values of *V*_g_, *V*_s_, total time in growth, and total time in shortening also differ. However, when these measurements are used in evaluating the *J* equation, the results for *J* are essentially the same (compare *J* across the four panels). This concurrence occurs because of the following:

- *V*_g_*(total time in growth) gives the total length increase that occurs during detected growth;
- *V*_s_*(total time in shortening) gives the total length decrease that occurs during detected shortening;
- *V*_g_*(time in growth) and *V*_s_*(time in shortening) depend on what is categorized as growth or shortening, but their sum *V*_g_*(time in growth) + *V*_s_*(time in shortening) yields the overall length change regardless of whether any particular segment was classified as growth or shortening.

### 2.4 Calculating *J* from the DI parameters

As indicated above, Equations 1 through 5 can work even if there is time in phases during which the MT length does not change (e.g., if a pause occurs or if a MT seed is empty for some amount of time). The remaining equations in this paper will depend on the three simplifying assumptions listed below, in order to obtain forms of the *J* equation that are common in the literature (e.g., (Hill & Chen, 1984; Walker et al., 1988; Dogterom & Leibler, 1993)). However, as will be discussed below, there is later experimental evidence indicating that physical MTs may deviate from some of these assumptions (Tran, Walker, & Salmon, 1997; Jánosi, Chrétien, & Flyvbjerg, 2002; Odde, Cassimeris, & Buettner, 1995; Gardner, Zanic, Gell, Bormuth, & Howard, 2011), which could cause complications when applying the equations.

Let

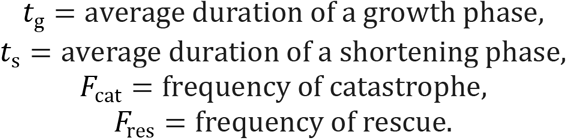

In our analysis, we calculate *F*_cat_ as (number of catastrophes)/(total time in growth) and *F*_res_ as (number of rescues)/(total time in shortening).

*Assumption* (i): Individual microtubule assembly/disassembly behavior is purely a two-state process where the two states are growth and shortening.

*Assumption* (ii): *t*_g_ = 1 /*F*_cat_.

*Assumption* (iii): *t*_s_ = 1 /*F*_res_.

#### 2.4.1 Derivation of *J*_AverageDuration_ using Assumption (i)

Under the two-state assumption (Assumption (i)), *P*_growth_ = *t*_g_/(*t*_g_ + *t*_s_) and *P*_shortening_ = *t*_s_/(*t*_g_ + *t*_s_). Substituting these formulas for *P*_growth_ and *P*_shortening_ into Equation 1 leads to

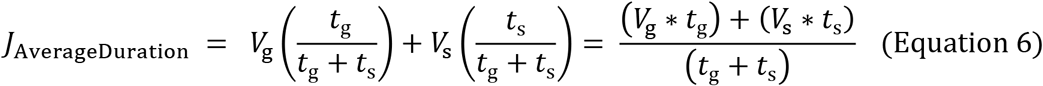

(Walker et al., 1988; Gliksman et al., 1992; Vorobjev & Maly, 2008).

#### 2.4.2 Derivation of *J*_DI_ using Assumpt ions (i), (ii), (iii)

*P*_growth_ and *P*_shortening_ can be calculated from the frequencies of catastrophe and rescue, if Assumptions (ii) and (iii) (Hill & Chen, 1984; Walker et al., 1988) are satisfied in addition to Assumption (i) above.

Under the Assumptions (i), (ii), and (iii),

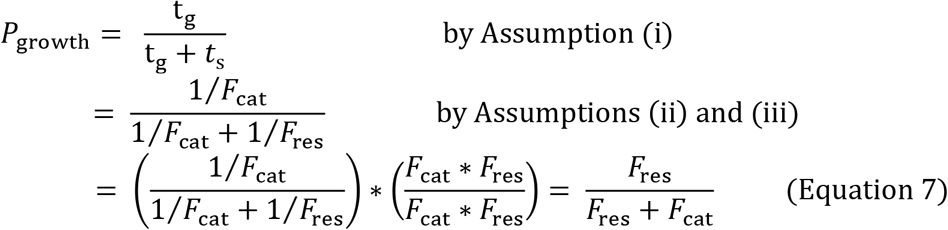

and

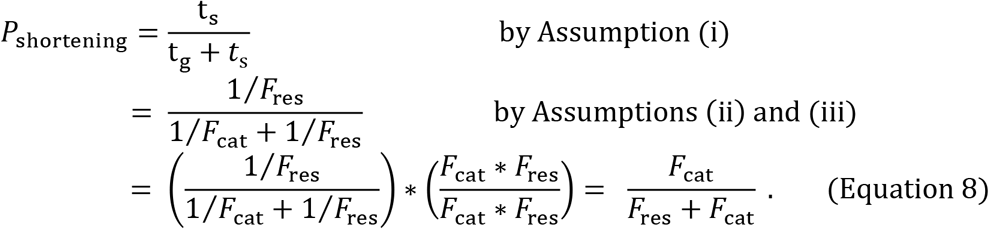

Substituting Equations 7 and 8 into Equation 1 yields

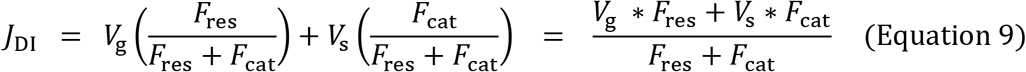

(Hill & Chen, 1984; Walker et al., 1988; Verde et al., 1992; Dogterom & Leibler, 1993; Maly, 2002). Maly also presents a drift coefficient equation (i.e., a *J* equation) that incorporates pauses, in addition to growth and shortening (Maly, 2002).

For the dilution simulations, Figure 9 compares the results of the *J*_DI_ equation and *J* measured directly from the net rate of change in average MT length, *J*_Net_, plotted as functions of post-dilution [free tubulin] (see also **Supplemental Figure S1**). The simulation results show that the *J*_DI_ equation fits the *J*_Net_ data fairly well, but that some deviation occurs in the [free tubulin] range from approximately 8 to 11 µM. This deviation in the intermediate range of [free tubulin] decreases with time after the dilution (Figure 9, compare **A** to **C** and **B** to **D**), but performing the measurements at such late times in physical experiments may be affected by complete depolymerizations (as occurred in Figure 9C at low [free tubulin]; further examined in Section 2.6.2).

**Figure 9:**
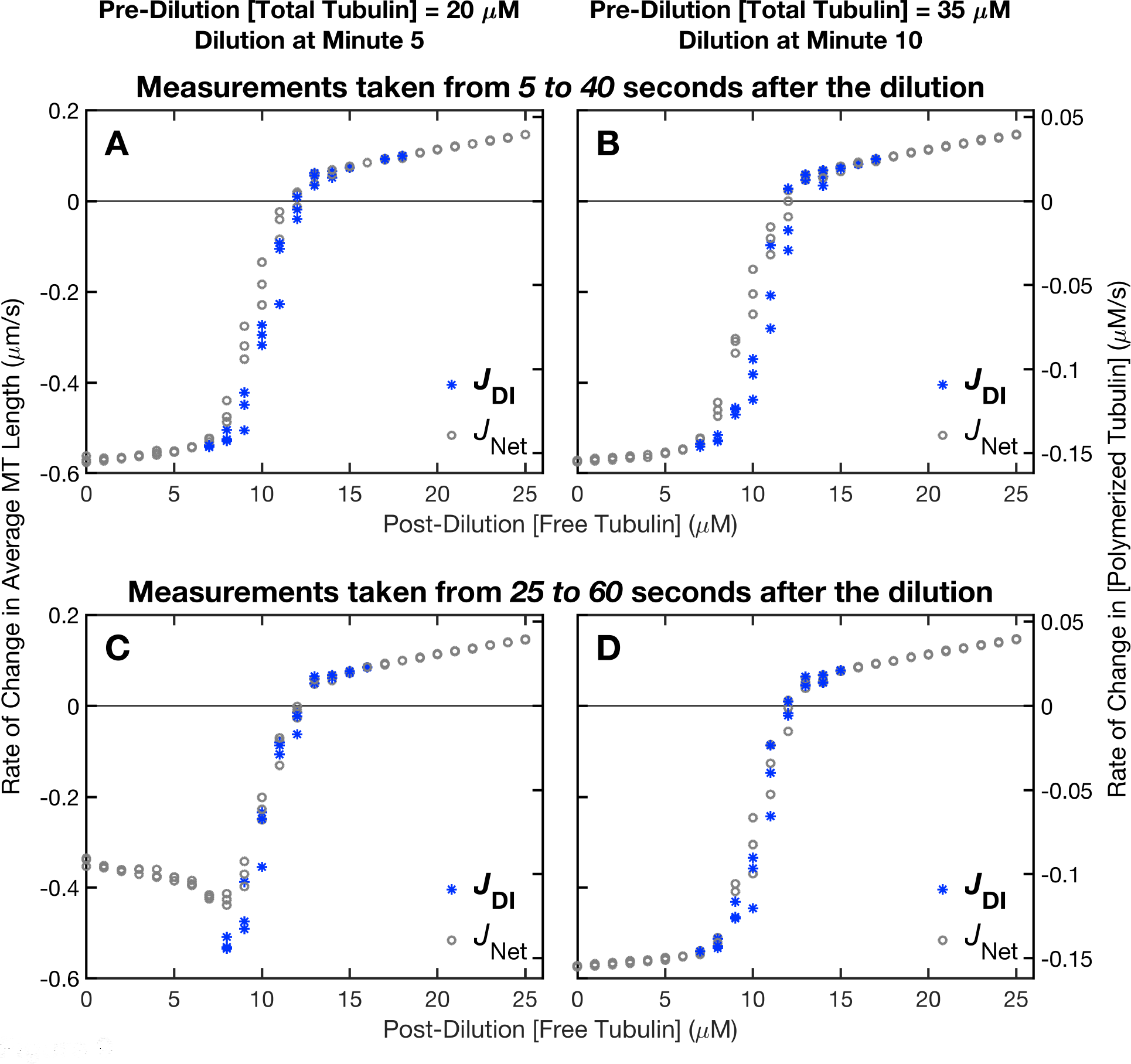
Comparison of the *J*_DI_ equation and the *J*_Net_ data in the dilution simulations. This figure shows a comparison of the results of the *J*_DI_ equation (asterisk symbols; Equation 9) to *J*_Net_ (circle symbols) for the dilution simulations. **(*A,C*)** Pre-dilution [total tubulin] = 20 µM, dilution time = minute 5. These conditions were chosen to be experimentally realistic. **(*B,D*)** Pre-dilution [total tubulin] = 35 µM, dilution time = minute 10. These conditions were chosen to produce very long microtubules, so that the mathematical equations could be tested over longer measurement periods without any complete depolymerizations (however, in experiments, problems such as spontaneous nucleation could arise at such high concentrations). ***Methods***: In each panel, *J*_Net_ and *J*_DI_ were evaluated over the same time period as each other: 5 to 40 s after the dilution in panels **A-B**; and 25 to 60 s after the dilution in panels **C-D**. The *J*_DI_ equation, *J*_DI_ = (*V*_g_ ∗ *F*_res_ + *V*_s_ ∗ *F*_cat_)/(*F*_res_ + *F*_cat_), was evaluated with measurements of the DI parameters (*V*_g_, *V*_s_, *F*_cat_, *F*_res_) obtained using the DI analysis method (Methods, Section 3.2.2). The *J*_Net_ data in **panel A** are re-plotted from Figure 4A. *J*_Net_ data points are plotted for each of the three independent replicates of the simulations at each value of post-dilution [free tubulin]. *J*_DI_ is plotted only when time in growth and time in shortening are non-zero (so that *F*_res_ and *F*_cat_ are both defined); for additional analysis, see **Supplemental Figure S1**. ***Interpretations***: *Comparison of J_DI_ and J*_*Net*_: The results of the *J*_DI_ equation evaluated with the DI measurements are close to but do not exactly match the *J*_Net_ data. In the range from ~8 to ~11 µM, *J*_DI_ underestimates *J*_Net_ when the measurements are taken shortly after the dilution (panels **A-B**). This discrepancy is largest at ~ 9 – 10 µM, which is the concentration range where *P*_growth_ and *P*_shortening_ are closest to being equal (Figure 4B), i.e., where DI is most robust. The underestimate does not occur when the measurements are taken later in time (panels **C-D**), but other problems may occur due to complete depolymerizations. *Effect of complete depolymerizations*: For pre-dilution [total tubulin] = 20 µM, comparison of *J*_Net_ in panels **A** and **C** shows that the lower arm of *J* is shifted upwards when the measurements are performed later in time. As will be examined further in Section 2.6.2, this shift is due to complete depolymerizations during the measurement period. In contrast, for pre-dilution [total tubulin] = 35 µM, where the MTs are longer at the time of the dilution, no shift occurs (compare *J*_Net_ in panels **B** and **D**).

#### 2.4.3 Complications in applying the *J*_DI_ equation to experimental systems

Recall that the derivation of the *J*_DI_ equation (Equation 9) depends on the simplifying Assumptions (i), (ii), and (iii) listed above. However, there is experimental evidence that physical microtubule behavior may deviate from these assumptions. This could cause complications in measuring the DI parameters to input into the equation.

Assumption (i) presumes that MTs do not exhibit any additional states besides growth and shortening. However, for example, (Tran et al., 1997; Jánosi et al., 2002) indicate the existence of a third state that is intermediate between growth and shortening. Detection of intermediate states in our simulations will be investigated in future work. For the analysis in this paper, we divided the length histories of individual MTs into only growth, shortening, or empty seed phases (the empty seed state is relevant when complete depolymerizations occur, as will be examined in Section 2.5).

The simplest scenario in which Assumptions (ii) and (iii) would hold is if the transition from growth to shortening is a first-order process with transition rate constant *F*_cat_, and the transition from shortening to growth is a first-order process with transition rate constant *F*_res_. Here, first-order means that the times until catastrophe for growing MTs and the times until rescue for shortening MTs are each exponentially distributed. In this case, the overall rate of catastrophe for the population is (*F*_cat_)*(# of growing MTs) and overall rate of rescue is (*F*_res_)*(# of shortening MTs).

However, there is evidence that times until catastrophe are not exponentially distributed, but instead follow a gamma distribution due to age-dependent catastrophe (e.g., (Odde et al., 1995; Gardner et al., 2011; Coombes, Yamamoto, Kenzie, Odde, & Gardner, 2013)). In this case, *F*_cat_ would be time-dependent; specifically the value of *F*_cat_ would increase over time during a growth phase (Gardner et al., 2011).

In our analysis, we calculated *F*_cat_ as (number of catastrophes)/(total time in growth). This provides an average *F*_cat_ value for the time period during which measurements were taken. If *F*_cat_ is age-dependent and the measurements were only taken early during growth phases, then the average *F*_cat_ would be underestimated.

### 2.5 Effect of complete depolymerizations

One situation in which Assumption (iii) fails is if MTs completely depolymerize. In this case, the transition from shortening to growth can occur by way of complete deploymerization followed by regrowth from the stable MT seed, rather than occurring only through rescue. Then, the transition frequency from shortening to growth would not be simply *F*_res_, and Equations 7 to 9 would not hold. If there is time between the end of complete deploymerization and the start of re-growth, then Assumption (i) and Equation 6 also break down.

One case where complete depolymerizations occur is if [free tubulin] is in the range where *J* = 0 in the constant [free tubulin] simulations (Figures 4C-D and 5C-D) (Jonasson et al., 2019). The [free tubulin] at which the transition from *J* = 0 to *J* > 0 occurs in the constant [free tubulin] simulations (Figure 4C, diamonds) is CC_NetAssembly_ (Figure 2, Table 1). CC_NetAssembly_ is also the [free tubulin] at which the transition from *J* < 0 to *J* > 0 occurs in the dilution simulation (Figure 4A, circles).

As would be expected, Figure 10 shows that the *J*_DI_ equation does not fit the *J*_Net_ data from the constant [free tubulin] simulations in the [free tubulin] range where *J* = 0 (i.e., [free tubulin] below CC_NetAssembly_).

**Figure 10:**
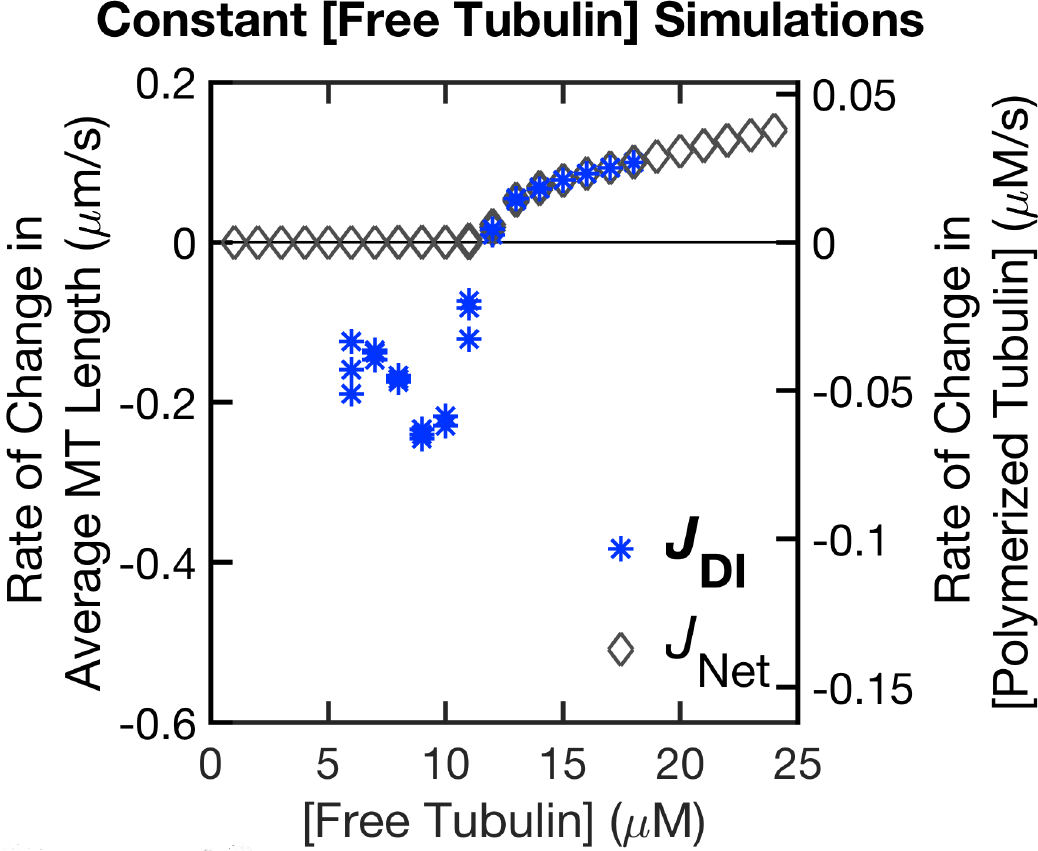
Comparison of the *J*_DI_ equation to the *J*_Net_ data in the constant [free tubulin] simulations. This figure shows a comparison of the *J*_DI_ equation (asterisk symbols; Equation 9) to *J*_Net_ (diamond symbols) for the constant [free tubulin] simulations. ***Methods***: The *J*_Net_ data are re-plotted from Figure 4C. The *J*_DI_ equation, *J*_DI_ = (*V*_g_ ∗ *F*_res_ + *V*_s_ ∗ *F*_cat_)/(*F*_res_ + *F*_cat_), was evaluated with measurements of the DI parameters (*V*_g_, *V*_s_, *F*_cat_, *F*_res_) obtained using the DI analysis method (Methods, Section 3.2.2). The DI measurements were taken during the same time period as the *J*_Net_ measurements. *J*_Net_ data points are plotted for each of the three independent replicates of the simulations at each value of [free tubulin]. *J*_DI_ is plotted only when time in growth and time in shortening are non-zero (so that *F*_res_ and *F*_cat_ are both defined). ***Interpretations***: The results of the *J*_DI_ equation evaluated with the DI measurements match *J*_Net_ well in the [free tubulin] range where *J*_Net_ > 0, but not in the [free tubulin] range where *J*_Net_ ≈ 0. In the range where *J*_Net_ ≈ 0, the MTs undergo complete depolymerizations back to seed; therefore, the rate of entering into growth is not simply *F*_res_; as discussed further in the main text, this situation violates assumptions used in deriving the *J*_DI_ equation. Instead, the population behavior is better described by *J*_DI_piecewise_ (Equation 10): *J*_DI_piecewise_ equals *J*_DI_ in the range where *J*_DI_ ≈ *J*_Net_ > 0 (“unbounded growth” regime), and *J*_DI_piecewise_ equals 0 in the range where *J*_Net_ ≈ 0 (“bounded growth” regime).

For constant [free tubulin] < CC_NetAssembly_, the average MT length will reach a steady-state value over time and the rate of change in average MT length will then be zero (*J* = 0) (Figures 4C and 5C). Thus, for constant [free tubulin] experiments or simulations, the *J*_DI_ equation (Equation 9) does not hold below CC_NetAssembly_ (Figure 10). Instead,

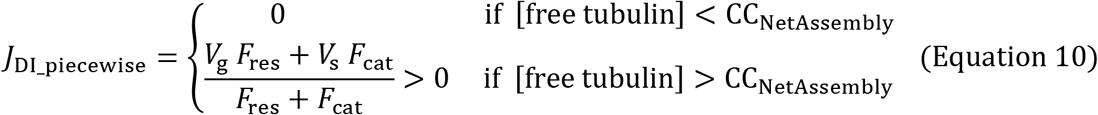

(Dogterom & Leibler, 1993). In the terminology of Dogterom et al., MTs exhibit “bounded growth” when *J* = 0 and “unbounded growth” when *J* > 0 (Dogterom & Leibler, 1993).

Depending on the specific application, empty and non-empty seeds may be considered separately; for example, in a system at polymer-mass steady state, *J* would be zero for the overall population of seeds, but would be positive for the empty seeds (since they cannot have shortening) and negative for the non-empty seeds (Vorobjev & Maly, 2008). In this case, Equations 6 to 9 could be applied to the population of non-empty seeds.

### 2.6 Timing of experimental steps and measurements in dilution systems

The accuracy of the measurement of *J* in dilution experiments can be affected by the timing of experimental steps.

#### 2.6.1 Delay for GTP cap to adjust to post-dilution [free tubulin]

After the time of the dilution, a delay before the start of the measurement period allows the GTP cap to adjust to the post-dilution [free tubulin]. Without the delay, *J* would be misestimated relative to its steady-state value, particularly at low values of [free tubulin] (Figure 11, **Supplemental Figure S2**).

**Figure 11:**
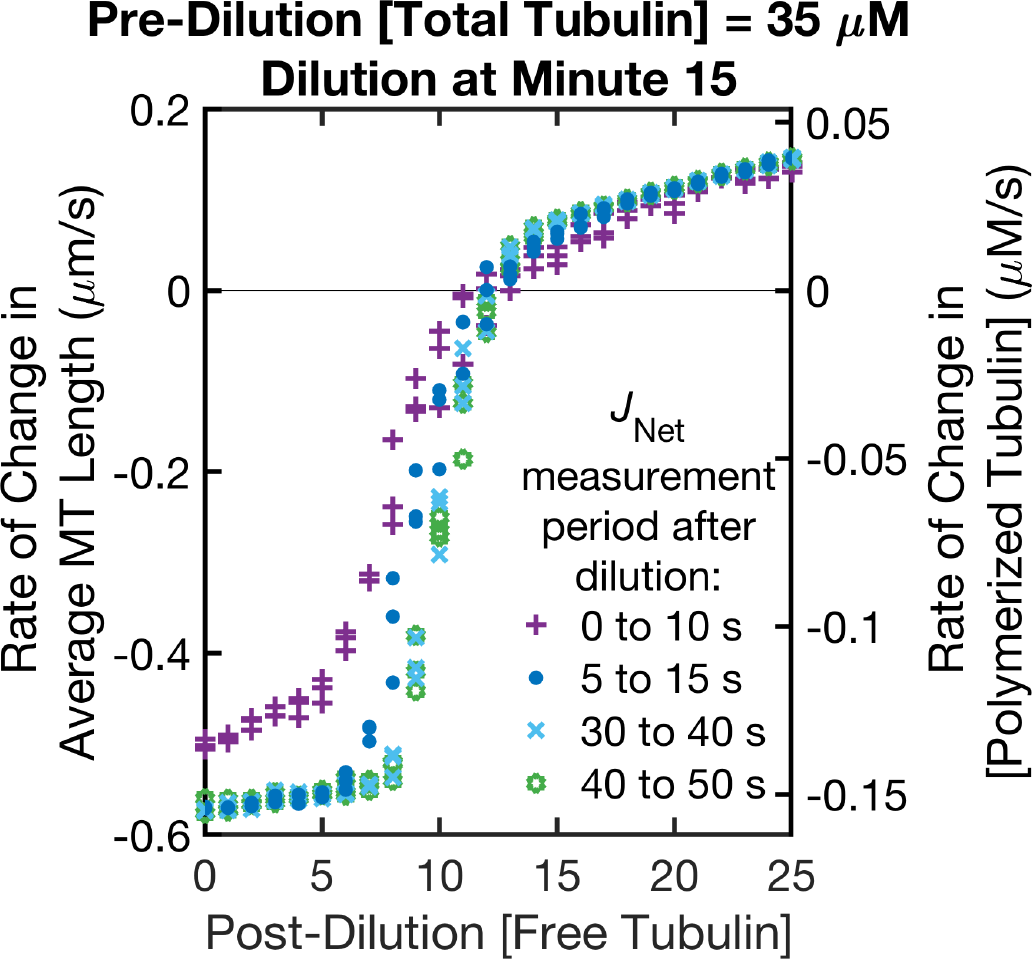
Effect on *J*_Net_ of varying the start time of the measurement period after the dilution. Measurements of *J*_Net_ are shown for 10-second intervals with varying start times after the dilution, as indicated in the key on the plot. Here, pre-dilution [total tubulin] = 35 µM, dilution time = minute 15. Analogous plots for varied pre-dilution [total tubulin] and dilution times are shown in **Supplemental Figure S2**. ***Methods***: *J* is calculated from the net rate of change in average MT length, *J*_Net_, over each of the 10-second intervals. Data points are plotted for each of three independent replicates of the simulations at each value of post-dilution [free tubulin]. ***Interpretations***: If the measurement period is started immediately after the dilution (0 - 10 s data, + symbols), the magnitude of *J* is underestimated relative to its steady-state value for all values of [free tubulin]. At very low and very high [free tubulin], a 5-second delay before the measurement period is sufficient for *J* to reach its steady-state value for the post-dilution [free tubulin]. For intermediate [free tubulin], it takes longer for *J* to reach its steady-state value (compare 5 - 15 s data to 30 - 40 and 40 - 50 s data). The need for a delay can be explained by evidence that the GTP cap requires time after the dilution to adjust to the new value of [free tubulin] (Duellberg, Cade, Holmes, & Surrey, 2016). Our simulation results shown here suggest that this adjustment takes longer in the intermediate range of [free tubulin], which is where growth and shortening are both occurring.

#### 2.6.2 Effect of complete depolymerizations on *J*

Ideally, dilution experiments should be performed so that measurements of *J* after the dilution can be taken before any MTs have completely depolymerized to the seed. If MTs that are too short are present at the time of the dilution, they will completely depolymerize during the measurement period, causing the lower arm of *J* to shift upwards. The lengths of the MTs during the measurement period can be affected by various factors of the experimental setup (e.g., pre-dilution [total tubulin], number of seeds) and timing of the experimental steps (e.g., time of dilution, start time of the measurement period, time duration of the measurement period). To test the effects of these factors on *J*, we performed simulations with different values of pre-dilution [total tubulin] and different times of dilution. For each, we measured *J*_Net_ over varying times periods. The results in Figures 12, 13, and 14 show that there are several different factors that each increase the number of complete depolymerizations that occur during the measurement period and that all can cause the lower arm of J to shift upwards:

**Figure 12:**
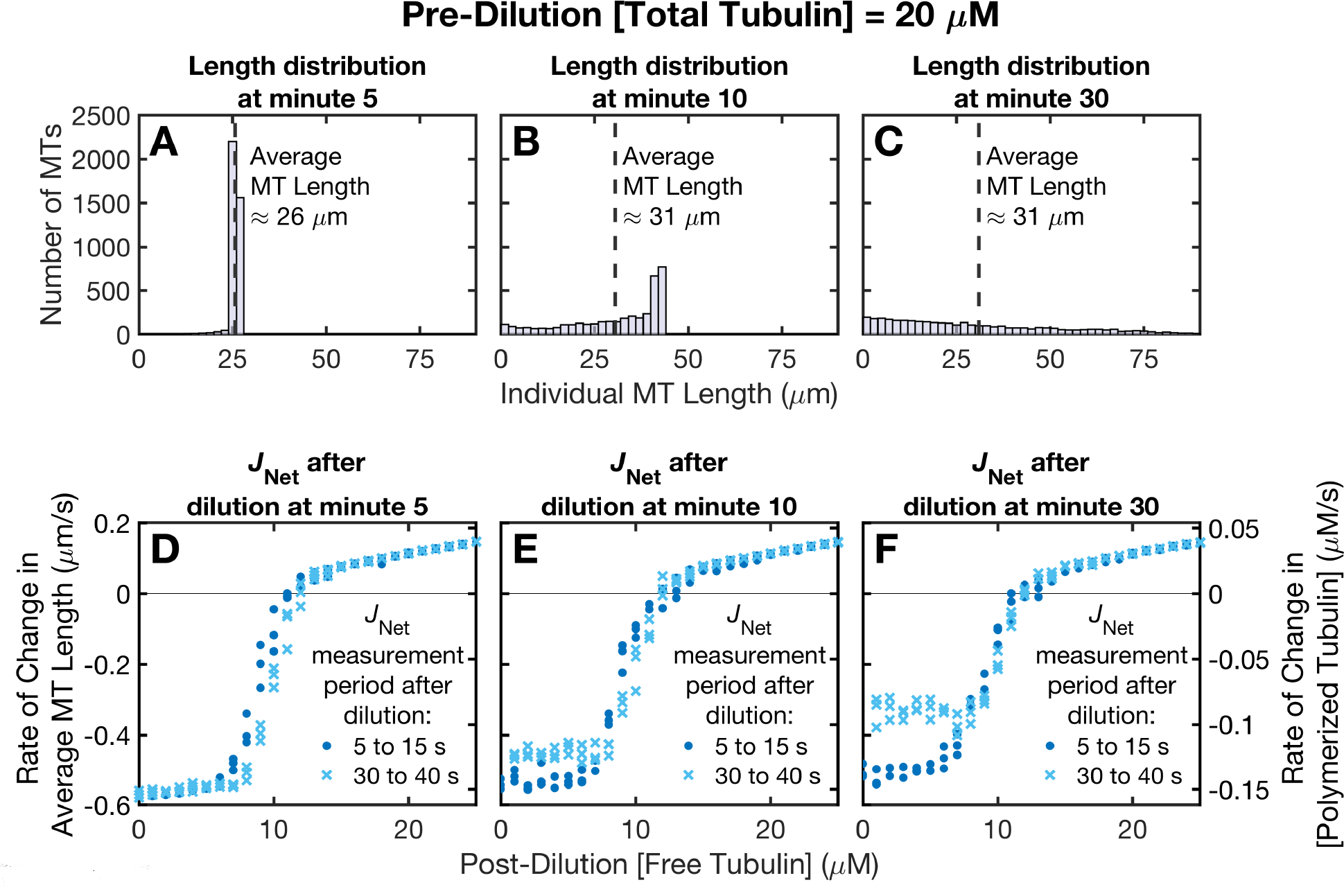
Effect on *J*_Net_ of varying the time at which the dilution is performed. Pre-dilution [total tubulin] = 20 µM. **(*A,B,C*)** Histograms of MT lengths at the time of the dilution: minute 5 (panel **A**), minute 10 (panel **B**), and minute 30 (panel **C**) of the simulation. Note that panel **A** is before the system has reached polymer-mass steady state and that panels **B-C** are after the system has reached polymer-mass steady state, which is when the average MT length is no longer changing over time but the length distribution can still change. **(*D,E,F*)** *J*_Net_ measured from 5 to 15 seconds after the dilution and 30 to 40 seconds after the dilution (as indicated in the keys on the plots), for each of the three different dilution times: minute 5 (panel **D**), minute 10 (panel **E**), and minute 30 (panel **F**). ***Methods***: In panels **A-C**, the three independent replicates of the simulations are aggregated in each histogram. For each post-dilution [free tubulin], the conditions up to the time of dilution are the same. Therefore, for the histograms, we aggregated the MTs that will be diluted into the different post-dilution [free tubulin]: *N* = 3900 = (50 MT seeds per post-dilution [free tubulin] per replicate) * (26 values of post-dilution [free tubulin]) * (3 replicates). In panels **D-F**, data points are plotted for each of three independent replicates of the simulations at each value of post-dilution [free tubulin]. **Interpretations:** The later the dilution time (i.e., the longer the amount of time that the MTs are allowed to compete before the dilution is performed), the more short MTs are present at the time of the dilution (compare across panels **A-C**). The shorter the MTs are at the time of the dilution, the sooner they undergo complete depolymerization after the dilution for low post-dilution [free tubulin] (see Figure 13). These complete depolymerizations cause the lower arm of *J* to shift upwards (compare across panels **D-F**).

**Figure 13:**
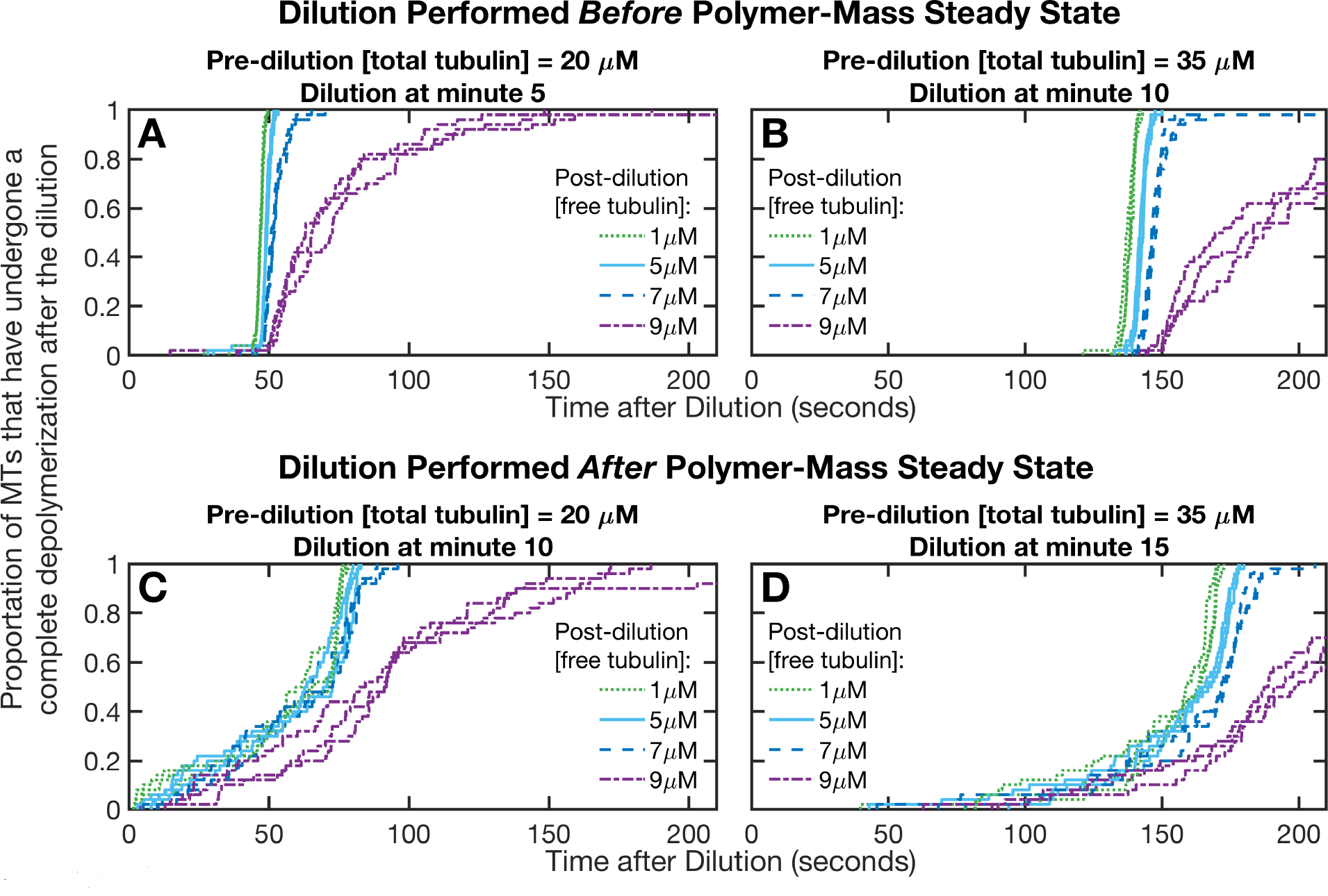
Effect of pre-dilution [total tubulin] and dilution time on the time until complete depolymerization after the dilution. This figure plots the proportion of MTs that have undergone a complete depolymerization after the dilution versus time after dilution. The dilution was performed either before (panels **A-B**) or soon after (panels **C-D**) polymer-mass steady state (Table 1) was reached in the pre-dilution competing systems. **(*A,C*)** Pre-dilution [total tubulin] = 20 µM, with either time of dilution = minute 5 of the simulation (panel **A**), or time of dilution = minute 10 (panel **C**). **(*B,D*)** Pre-dilution [total tubulin] = 35 µM, with either time of dilution = minute 10 (panel **B**), or time of dilution = minute 15 (panel **D**). ***Methods***: The time of complete depolymerization of each MT was measured as the time at which the MT length first decreased to within 2 subunit lengths above the seed. In each panel, curves are plotted for each of three independent replicates of the simulations at each value of post-dilution [free tubulin] shown. ***Interpretations***: When the dilution is performed before polymer-mass steady state (panels **A-B**), all the MTs in the population have similar lengths at the time of the dilution (e.g., Figure 12A). In this case, and if post-dilution [free tubulin] is low (panels **A,B**, data series for 1, 5, 7 µM post-dilution [free tubulin]), then most of the MTs reach complete depolymerization around the same time point after the dilution. In contrast, when the dilution is performed after polymer-mass steady state (panels **C-D**), the range of MT lengths at the dilution time is more spread out and there are more short MTs (e.g., Figure 12B-C). In this case, more of the MTs experience complete depolymerization sooner after the dilution time (compare panels **A-B** to panels **C-D**). Thus, if the dilution is performed before polymer-mass steady state, then, after the dilution, there will be a longer time period during which *J* measurements can be taken before any MTs completely depolymerize. Additionally, when the pre-dilution [total tubulin] is higher, complete depolymerizations do not begin occurring until later in time after the dilution (compare left and right columns).

**Figure 14:**
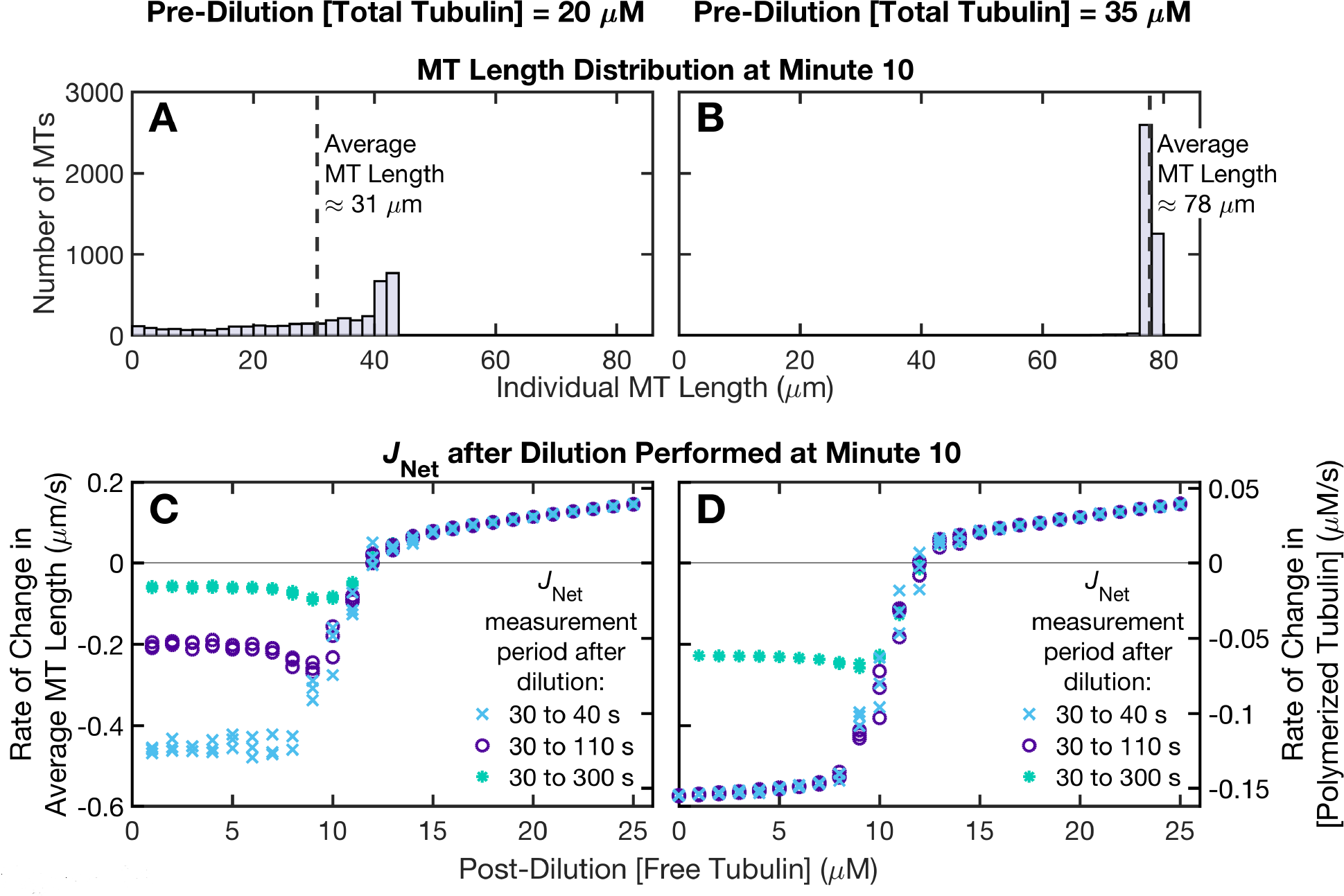
Effects on *J*_Net_ of changing the value of pre-dilution [total tubulin] and the duration of the measurement period. Left column: pre-dilution [total tubulin] = 20 µM; right column: pre-dilution [total tubulin] = 35 µM. Time of dilution = minute 10 of the simulation. **(*A,B*)** Histograms of MT lengths at minute 10. **(*C,D*)** *J*_Net_ measured over time periods of varying durations, all starting 30 seconds after the time point of dilution. ***Methods***: The histograms in panels **A-B** are aggregated as described in the Figure 12 legend (*N* = 3900). Data points in panels **C-D** are plotted for each of three independent replicates of the simulations at each value of post-dilution [free tubulin]. Note that panel **A** of this figure is the same as panel **B** of Figure 12, and that the 30 to 40 s data series (x symbols) in panel **C** of this figure is the same as the 30 to 40 s data series in panel **E** of Figure 12. **Interpretations:** When the duration of the measurement time period is longer, more complete depolymerizations occur (Figure 13), causing the lower arm of *J* to shift upwards (compare data series within each of panels **C-D**). This upward shift is greatest at the lowest values of post-dilution [free tubulin]. At the time point of the dilution, the MTs are longer for the 35 µM pre-dilution [total tubulin] (panel **B**) than for the 20 µM pre-dilution [total tubulin] (panel **A**). Hence, the measurements can be performed over a longer time period for the 35 µM than for the 20 µM before the effect of complete depolymerizations on the shape of *J* is observed (compare panel **D** to panel **C**). However, as noted earlier, performing experiments at 35 µM may be unrealistic because of problems as such spontaneous nucleation.

- later of time of dilution (compare across the three columns in Figure 12);
- later start time of the measurement period (compare the two data series within each of panels **D,E,F** in Figure 12);
- lower pre-dilution [total tubulin] (compare panels **A,C** to **B,D** within each of Figures 13 and 14);
- longer duration of the measurement period (compare the three data series within each of panels **C,D** in Figure 14).

Traditionally, dilution experiments are performed by growing MTs to polymer-mass steady state under competing conditions (constant [tubulin total]) and then transferring the MTs to new concentrations of [free tubulin]. However, as shown in Figure 15, some MTs in a population will begin to have noticeable shortening phases even before the pre-dilution competing system has reached polymer-mass steady state. Furthermore, the longer the system is allowed to run after reaching polymer-mass steady state, the more MTs will have undergone complete depolymerization (Figure 12). In contrast, if the dilution is performed before the competing system has reach polymer-mass steady state, this can increase the time period after the dilution before any MTs completely depolymerize (Figure 13).

**Figure 15:**
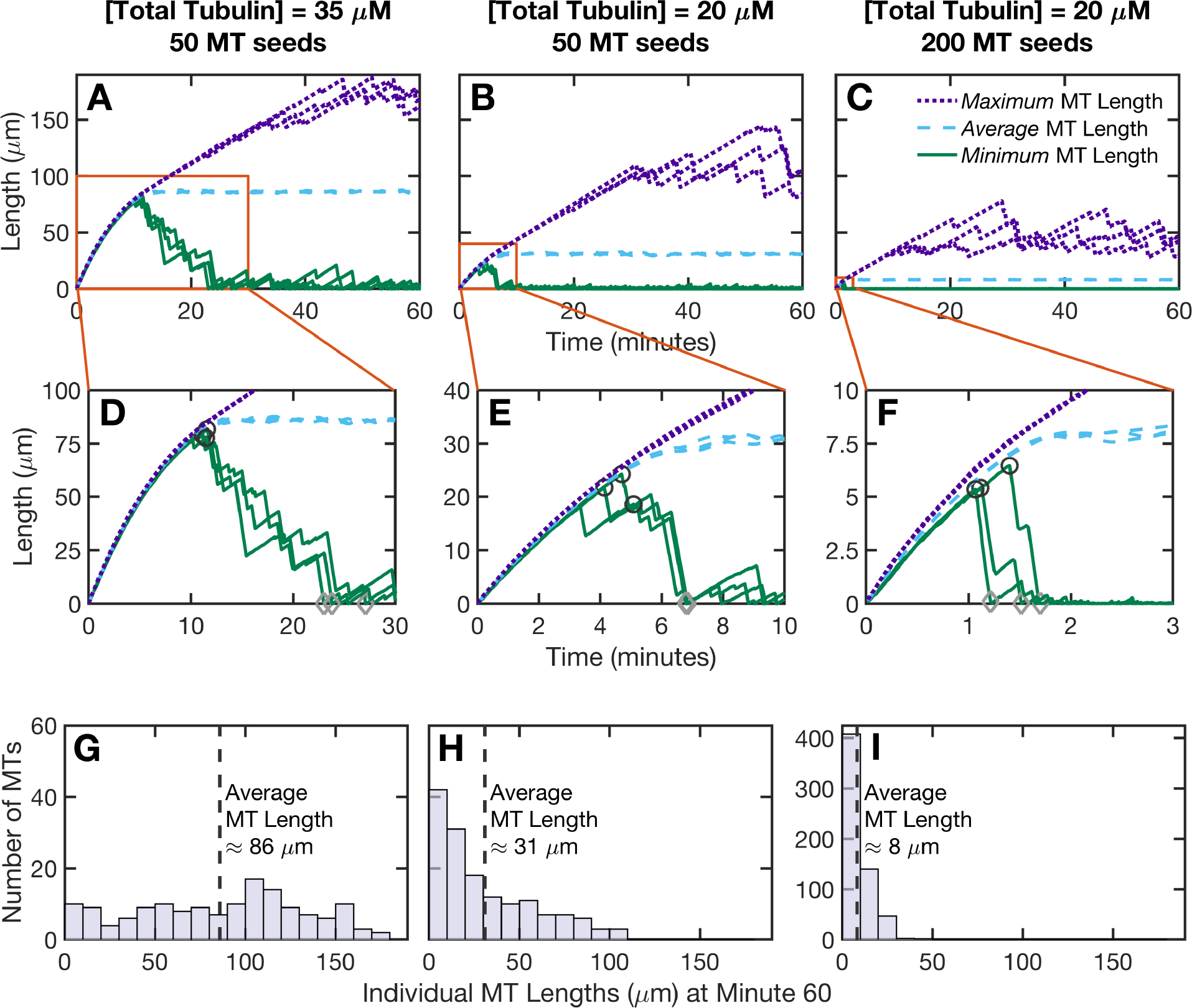
Minimum, average, and maximum MT length versus time for competing simulations. **(*A-C*)** The minimum (solid lines), average (dashed lines), and maximum (dotted lines) MT length of a competing population versus time. **(*D-F*)** Zooms-in of the indicated regions of panels **A-C**. The data in panels **A-F** are plotted to examine the range of lengths present in competing populations at potential dilution times. In panels **D-F**, the circle symbols indicate the point when the minimum MT length is at its highest value, and the diamond symbols indicate the point when the minimum MT length first decreases to within 2 subunit lengths of the seed. **(*G-I*)** Histograms of the MT length distribution at minute 60 of the simulations. *First column (panels **A,D,G**)*: [Total tubulin] = 35 µM, number of stable MT seeds = 50. *Second column (panels **B,E,H**)*: [Total tubulin] = 20 µM, number of stable MT seeds = 50. *Third column (panels **C,F,I**)*: [Total tubulin] = 20 µM, number of stable MT seeds = 200. ***Methods***: In panels **A-F**, curves are plotted for each of three independent replicates of the competing simulations. In panels **G-I**, the three replicates are aggregated in the histograms. In panels **G-H**, *N* = 150 = (50 MT seeds per replicate) × (3 replicates). In panel **I**, *N* = 600 = (200 MT seeds per replicate) × (3 replicates). ***Interpretations:**Behaviors in competing systems*: Initially, in a competing system, [free tubulin] equals [total tubulin]; over time MTs grow and take up free tubulin, so [free tubulin] decreases and [polymerized tubulin] increases until both level off when polymer-mass steady state is reached (e.g., see Figure S1 of (Jonasson et al., 2019)). Early in time, [free tubulin] is high, so all the MTs in the population are growing; thus, in panels **A-F**, the minimum (solid lines), average (dashed lines), and maximum length (dotted lines) are close to each other. Over time, [free tubulin] decreases, and some MTs start shortening (solid lines). Polymer-mass steady state is when the average MT length (dashed line) is no longer changing with time (other than small fluctuations around the average). If the system is run for a very long duration of time, then the length distribution eventually approaches an exponential-like distribution, meaning that there are many short MTs (e.g., histograms in panels **H-I**). *Relevance to choosing the optimal dilution time in a dilution experiment*: Traditionally, dilutions are performed sometime after the system has reached polymer-mass steady state (e.g., (Carlier et al., 1984a)). One might have expected that running the system for a longer duration of time after reaching polymer-mass steady state would produce longer MTs. Indeed, the maximum MT length (dotted lines) does keep increasing with time after polymer-mass steady state. However, the minimum MT length (solid lines) starts to decrease even before polymer-mass steady state. Performing the dilution before this length decrease begins (e.g., at time of circle symbol) will maximize the duration of time during which the *J* measurements can be performed before complete depolymerizations start occurring. The later the dilution is performed (e.g, after time of diamond symbol), the more MTs will completely depolymerize during the measurement period. If complete depolymerizations occur, then the lower arm of *J* will shift upwards (Figures 12 and 14). Also, note that the simulations in all prior figures were performed with 50 MT seeds. The right column (panels **C,F,I**) of this figure shows that increasing the number of MT seeds makes the MTs shorter and causes the minimum MT length to start decreasing sooner.

Thus, although one might think that it would be better to allow the system to run for a longer duration of time before the dilution, this is true only to a point. After some amount of time, MT lengths start to redistribute toward an exponential-like length distribution (e.g., Figure 15E-F; see also (Fygenson, Braun, & Libchaber, 1994)). If the dilution is performed later time, more MTs will be short and the lower arm of the measured *J* curve will increasingly be shifted upwards because of the impact of complete depolymerizations (Figure 12, compare progression from first column to last column).

### 2.7 Timing of measurements in constant [free tubulin] systems

As discussed above, measurements of *J* in dilution experiments are sensitive to the timing of experimental steps. Similarly, in constant [free tubulin] systems, *J* also depends on when the measurements are performed. Specifically, if measurements of *J* are performed too early in time, then *J* will be overestimated, particularly for [free tubulin] near CC_NetAssembly_ (the [free tubulin] at which the steady-state *J* transitions from being zero to being positive), as discussed in (Jonasson et al., 2019) and illustrated here in Figure 16. To obtain the correct steady-state value of *J*, the measurements should be performed after the system has reached polymer-mass steady state (for [free tubulin] < CC_NetAssembly_) or polymer-growth steady state (for [free tubulin] > CC_NetAssembly_). When using measurements of *J* to determine the value of CC_NetAssembly_, the measurements must be taken when *J* has reached its steady-state value; reaching this state will take longer the closer [free tubulin] is to CC_NetAssembly_.

**Figure 16:**
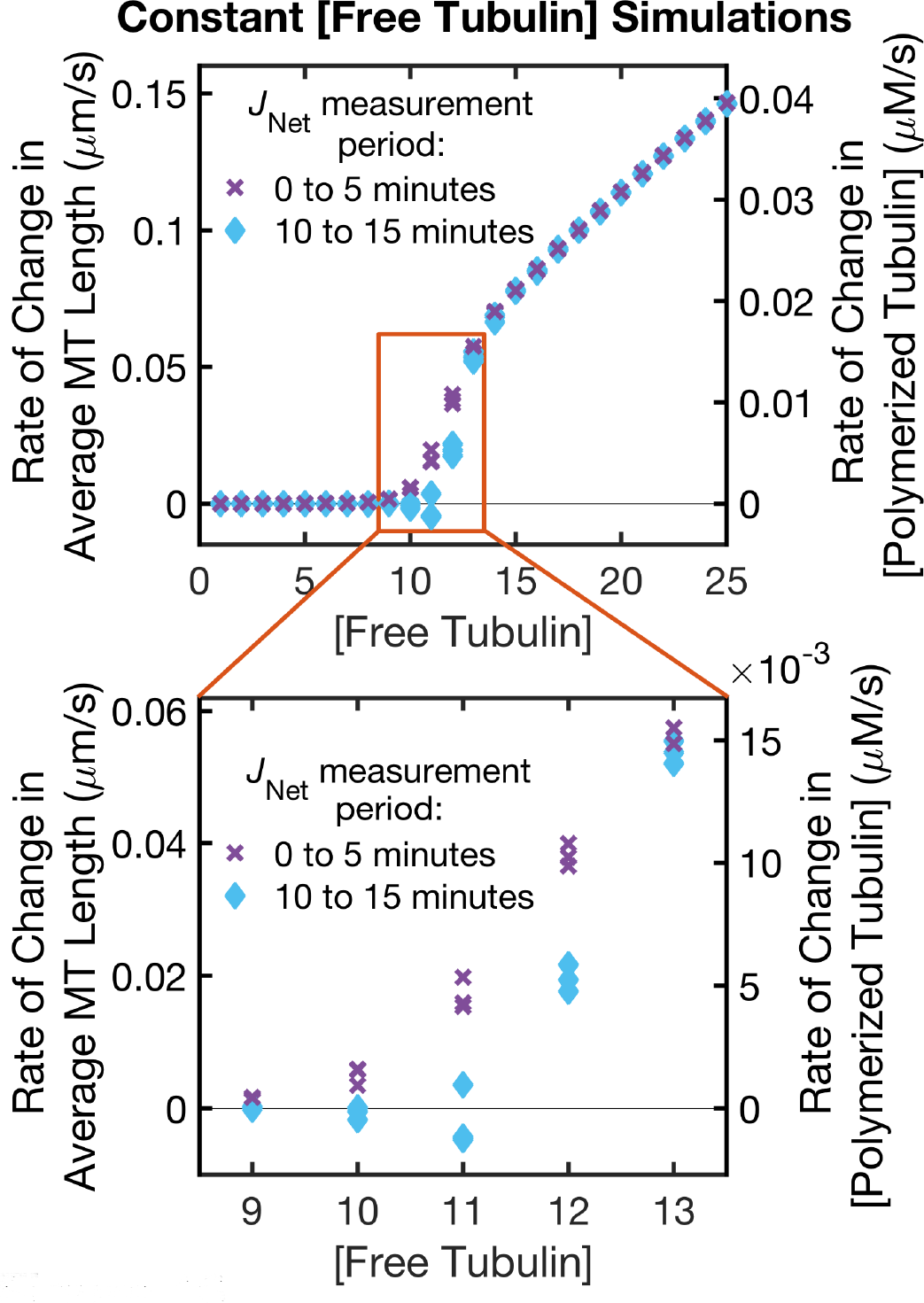
Effect of measurement start time on *J*_Net_ in the constant [free tubulin] simulations. For the constant [free tubulin] simulations, this figure shows *J*_Net_ calculated from 0 to 5 minutes (x symbols) and 10 to 15 minutes (diamond symbols). ***Methods:*** Data points are plotted for each of three independent replicates of the simulations at each value of [free tubulin] (same simulation runs as in earlier figures, e.g., Figures 4C-D, 5C-D). ***Interpretations:*** For [free tubulin] < CC_NetAssembly_ (CCs defined in Figure 2 and Table 1), the system reaches polymer-mass steady state where the average MT length (Figure 5C) is no longer changing with time (*J* = 0). For [free tubulin] > CC_NetAssembly_, the system reaches polymer-growth steady state where the average MT length (Figure 5C) increases at a rate that is constant with time (*J* > 0, with *J* at a constant value for each [free tubulin]). If the measurements of *J* are performed before the appropriate steady state is reached, then *J* will be overestimated (compare 0 - 5 minutes to 10 - 15 minutes). This overestimate will be most noticeable when [free tubulin] is near CC_NetAssembly_, because reaching steady state takes longer the closer [free tubulin] is to CC_NetAssembly_. See also relevant discussions in (Jonasson et al., 2019).

If measurements are performed when *J* has reached its steady-state value in dilution experiments and in constant [free tubulin] experiments, then *J* from the two types of experiments will be superimposed for [free tubulin] > CC_NetAssembly_ (see Figure 6C-D of (Jonasson et al., 2019)).

### 2.8 Summary and Practical Implications

#### 2.8.1 Various forms of the *J* equation relate individual and population dynamics

The *J* equation relates the flux of subunits into and out of polymer (or rate of change in average filament length) to growth and shortening behaviors of individual MTs. To our knowledge, versions of this equation were first presented by (Hill & Chen, 1984). Since then, varied forms of the *J* equation have appeared in the literature; these forms differ in attributes including the types of experimental data used as input. From looking at variants of the *J* equation (e.g., *J*_General_, *J*_Time_, *J*_TimeStep_, *J*_DI_), it might not be obvious how they relate to each other (i.e., how to convert between different forms). Additionally, even for the same version of the equation, different authors have used many different variable names for the quantities in the equation. We show how to algebraically convert between different forms of the equation and examine the assumptions needed for the forms to be equivalent. Specifically, the *J*_Time_ and *J*_TimeStep_ equations are algebraically equivalent to the *J*_DI_ equation if the following assumptions are met: (i) individual microtubule assembly/disassembly behavior is purely a two-state process where the two states are growth and shortening; (ii) the average duration of a growth phase equals 1/*F*_cat_; and (iii) the average duration of a shortening phase equals 1/*F*_res_.

By using the *J* equation, measurements on individual MTs (inputs into the equation, which vary among the different forms) can be used to calculate the population-level flux behavior (output of the equation). Since it is technically difficult to measure individual-level and population-level behaviors simultaneously in physical experiments, use of the *J* equation enables one to obtain information about both scales from measurements only at the individual scale. However, correct application of any form of the equation requires understanding the conditions under which that form holds and understanding how experimental design and execution can affect the measurements, which we illustrate with computational simulations of experiments.

#### 2.8.2 Comparing versions of the *J* Equation for their validity and usefulness

The variants of the *J* equation differ in the specific measurements on individuals that are used to evaluate the equation.

For example, some of the variants use dynamic instability parameters, obtained by following particular individuals for long periods of time. In particular, the *J*_Time_ equation (Equation 4) uses *V*_g_, *V*_s_, total time in growth, and total time in shortening. The *J*_DI_ equation (Equation 9) also uses *V*_g_ and V_s_, but calculates the probabilities of growth and shortening using *F*_cat_ and *F*_res_ instead of from the total times in growth and shortening (Hill & Chen, 1984; Walker et al., 1988; Verde et al., 1992; Dogterom & Leibler, 1993; Maly, 2002).

In contrast, the *J*_TimeStep_ equation (Equation 5), which is the drift coefficient formula of (Komarova et al., 2002), uses displacements of many individuals followed over short time steps and does not require the same individuals to be followed over long periods of time. Thus, experimentalists can choose the type of measurement that is most feasible for their experiments and then use the corresponding form of the *J* equation.

To assess the utility and accuracy of the above variants of the *J* equation under different conditions, we compared each to the value of *J*_Net_ as observed in our simulations. *J*_Net_ is the net rate of change in average MT length between two time points. For simulation data, *J*_Net_ provides the true net rate of change that occurs in any particular run of the simulation, and is therefore a useful baseline for comparisons.

As observed in the dilution simulations, the *J*_Time_ and *J*_TimeStep_ equations are less sensitive to the measurement time period than is the *J*_DI_ equation. Specifically, the results of both the *J*_Time_ and *J*_TimeStep_ equations closely match the *J*_Net_ data (Figures 6A and 7A). In contrast, the results of the *J*_DI_ equation deviate from the *J*_Net_ data if the measurements are taken too soon after the dilution (Figure 9, compare panels **A-B** to **C-D**). Additionally, if the *J*_DI_ measurement period is too short, few transitions will be detected, and the output of the *J*_DI_ equation will be very noisy.

As observed in the constant [free tubulin] simulations, the *J*_Time_ and *J*_TimeStep_ equations match the *J*_Net_ data even if there are complete depolymerizations (Figures 6B and 7B), but the *J*_DI_ equation does not (Figure 10). The *J*_DI_ equation uses the rescue frequency, *F*_res_, as the rate of transitioning from shortening to growth. When there are complete depolymerizations, transitions from shortening to growth can occur not only by rescue but also by re-growth from the MT seed following a complete depolymerization. The *J*_DI_ equation does not hold in this case, because the overall rate of transition from shortening to growth is greater than *F*_res_. Instead J_DI_piecewise_ holds (Equation 10; see also (Dogterom & Leibler, 1993)).

In both of the above cases, *J*_Time_ fits *J*_Net_ (Figure 6) better than *J*_DI_ fits *J*_Net_ (Figures 9, 10). Interestingly, the *J*_Time_ and *J*_DI_ equations both use measurements from DI analysis. The *J*_DI_ equation uses *F*_cat_ and *F*_res_, which are calculated from the numbers of catastrophes and rescues divided by the total times in growth and shortening, respectively. If the measurement period is not long enough to capture a significant number of transitions, then *F*_cat_ and *F*_res_ can be inaccurate and imprecise. The *J*_Time_ equation also uses total times in growth and shortening, but does not require knowing the number of transitions. Since *J*_Time_ fits the data better than *J*_DI_, *J*_Time_ may provide a more experimentally accurate way to determine *J*. More specifically, if one is performing DI analysis, then using the total time in growth and shortening directly in *J*_Time_ may provide a more accurate estimate of *J* than using total time in growth and shortening to calculate *F*_cat_ and *F*_res_ and then using *F*_cat_ and *F*_res_ in *J*_DI_.

As mentioned earlier, the DI analysis used in both in *J*_Time_ and *J*_DI_ requires following specific individuals over long times, whereas *J*_TimeStep_ uses the displacements of many individuals over short time steps and does not require following the same individuals for long periods of time. Moreover, *J*_TimeStep_ fits *J*_Net_ (Figure 7) as closely as *J*_Time_ fits *J*_Net_ (Figure 6). Thus, *J*_TimeStep_ may be more practical than *J*_Time_ or *J*_DI_.

*J*_Time_ and *J*_TimeStep_ work without needing to measure *F*_cat_ and *F*_res_. However, if one can obtain accurate measurements of *F*_cat_ and *F*_res_, then they provide information about the dynamicity of individual MTs within a population that is not provided by *J* itself (e.g., (Hill & Chen, 1984; Verde et al., 1992; Dogterom & Leibler, 1993)). Another quantity that also provides information about dynamicity is the “diffusion coefficient” of the MT lengths (e.g., (Hill, 1987; Verde et al., 1992; Dogterom & Leibler, 1993; Komarova et al., 2002; Vavylonis, Yang, & O’Shaughnessy, 2005; Mirny & Needleman, 2010)).

#### 2.8.3 Considerations specific to implementing dilution experiments

For dilution experiments, the accuracy of the measurement of *J* can be affected by various factors such as experimental set up (e.g., tubulin concentration, number of seeds) and timing of experimental steps (e.g., dilution time, measurement time period).

The MT lengths at the time of dilution depend on the pre-dilution [total tubulin], the number of seeds, and the point in time when the dilution is performed (Figures 12A-C, 14A-B, 15). If any MTs are too short at the time of dilution, complete depolymerizations occur soon after the dilution (Figure 13) and cause the lower arm of *J* to shift upwards (Figures 12D-F, 14C-D). To avoid short MTs, one might have expected that the dilution should be performed after the pre-dilution competing system has reached polymer-mass steady state and that waiting longer before the dilution would lead to longer MTs. However, as shown by Figures 12, 13, and 15 together, the ideal time to perform the dilution is before polymer-mass steady state because the proportion of short MTs in a population increases after this ideal time.

Additionally, to allow the GTP cap to adjust to the post-dilution [free tubulin] (Duellberg, Cade, Holmes, & Surrey, 2016), a delay is needed before beginning measurements after the dilution (Figures 11, **S2**). However, if the end of the measurement period is too late in time after dilution, complete depolymerizations will occur (Figures 12, 13, 14).

Thus, in designing a dilution experiment, it is necessary to account for both the requirement for a delay and the need to avoid complete depolymerizations during the measurement period. An experimental design that maximizes the length of the shortest microtubules in a population at the time of dilution will be most likely to lead to accurate measurements of *J*.

#### 2.8.4 Broader implications of the *J* equation for steady-state polymers

The *J* equation provides a way to understand how dynamic instability relates to the critical concentrations CC_Elongation_ and CC_NetAssembly_ (Figure 2). CC_Elongation_ is the [free subunit] above which *V*_g_ is positive, whereas CC_NetAssembly_ is the [free subunit] above which the steady-state value of *J* is positive. We have previously proposed that the separation between CC_Elongation_ and CC_NetAssembly_ may account for the behavioral differences between MTs and actin (Jonasson et al., 2019).

For polymer types that display instability, such as MTs, there are values of [free subunit] at which growth and shortening occur simultaneously within a population (i.e., *P*_shortening_ and *P*_growth_ are both positive). Then, as seen from *J*_general_ = *V*_g_ ∗ *P*_growth_ + *V*_s_ ∗ *P*_shortening_ (Equation 1), *V*_g_ and *J* will be different from each other (Figures 3,4). CC_Elongation_ and CC_NetAssembly_ will therefore be different from each other (Figure 2).

In contrast, for polymer types, such as actin, that do not display (detectable) dynamic instability, either *P*_shortening_ ≈ 1 or *P*_growth_ ≈ 1 at any particular subunit concentration. In other words, the population and individuals will have the same behavior, i.e., all individuals are growing or all individuals are shortening. In this case, *J* ≈ *V*_g_ whenever *J* > 0, and CC_Elongation_ and CC_NetAssembly_ would be the same or experimentally indistinguishable.

## 3 METHODS

### 3.1 Simulations

The simulations in this paper used the dimer-scale computational model of MT dynamics that was originally introduced in (Margolin et al., 2011; Margolin et al., 2012). Specifically, expect for minor changes in the amount of information being outputted, we used the same implementation of the simulation that was used in (Jonasson et al., 2019), therein referred to as the “detailed model”.

In this dimer-scale model, each MT is composed of 13 protofilaments, and each protofilament is a chain of discrete subunits representing tubulin dimers. The MT length is the average of the 13 protofilament lengths, with 1 subunit length equaling 8 nm. In the simulations, there is no physical boundary that would limit MT lengths.

The biochemical events in the model are attachment and detachment of subunits to/from protofilament tips, formation and breakage of lateral bonds between adjacent subunits in neighboring protofilaments, and hydrolysis of GTP-tubulin subunits to GDP-tubulin subunits. The kinetic rate constants for these processes are user-inputted values and depend on the nucleotide state of the subunits involved in each event. All attachment and detachment event occur at the tips of the protofilaments. One subunit can attach to a tip at a time, and any subunit or oligomer of subunits that is not laterally bonded to a neighboring protofilament can detach from a tip.

The length change behavior of an individual MT over time is an emergent property, resulting from the execution of the kinetic events described above. Consequently, the values of the DI parameters, [polymerized tubulin], and *J* are also emergent properties. Additionally, in competing systems, the value of [free tubulin] is another emergent property.

In this paper, all simulations were performed in a volume of 500 fL (= 5.00 × 10^−13^ L) with MTs growing from 50 stable non-hydrolyzable GTP-tubulin seeds, except in Figure 15C,F, which has 200 seeds. We used the kinetic rate constants from Parameter Set C of (Margolin et al., 2012), which was tuned to approximately match the plus-end dynamics of mammalian MTs at 10 µM as reported in (Walker et al., 1988). This parameter set was also used in (Jonasson et al., 2019).

Please see Table 1 for descriptions of the types of simulations: competing; non-competing or constant [free tubulin]; and dilution.

### 3.2 Analysis

#### 3.2.1 Calculation of *J*_Net_

The net rate of change in average MT length is determined from

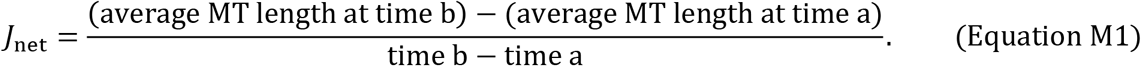

In the simulation outputs, the average MT length is the average of the individual MT lengths for all MT seeds in the population, and the length of each individual MT is the average of its 13 protofilament lengths.

Thus, the rate of change in average MT length can be converted to the rate of change in [polymerized tubulin] as follows:

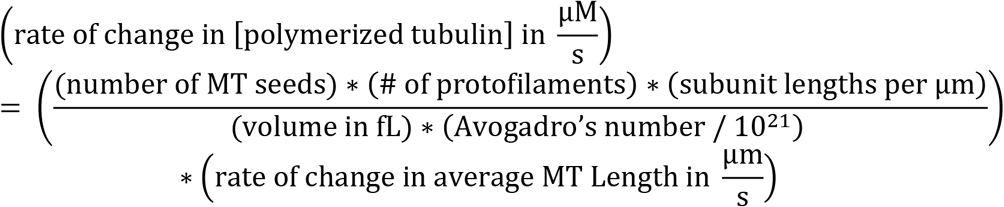

#### 3.2.2 DI analysis method

To identify growth and shortening phases in the MT length histories and to calculate the DI parameters, we use an automated DI analysis method (presented in the Supplemental Methods of (Jonasson et al., 2019)). Briefly, the DI analysis method identifies peaks and valleys in the data such that the length change between a peak and neighboring valley is greater than or equal to a user-defined threshold. For the analysis in this paper, we set the threshold to 25 subunit lengths (200 nm) to be comparable to detection limits in typical light microscopy. The ascent from a valley to a peak is classified as a growth phase and the descent from a peak to a valley is classified as a shortening phases. The DI parameters are calculated by

> *V*_g_ = total length change during growth phases / total time in growth phases,
>
> *V*_s_ = total length change during shortening phases / total time in shortening phases,
>
> *F*_cat_ = total number of catastrophes / total time in growth phases,
>
> *F*_res_ = total number of rescues / total time in shortening phases.

For more detailed information about the DI analysis method, please see the Supplemental Methods of (Jonasson et al., 2019).

#### 3.2.3 Time-step analysis method

The drift coefficient (*v*_d_) formula of (Komarova et al., 2002) provides the basis for the time-step analysis method used to evaluate the *J*_TimeStep_ = 𝑣_d_ = (∑*s*_i_)/(∑*t*_i_) (Equation 5). Here, we implemented the analysis as described in the Supplemental Methods of (Jonasson et al., 2019). Briefly, to apply the *J*_TimeStep_ equation to our simulation data, the length history of each MT was divided into 1-second time steps (*t*_i_), and the displacement (*s*_i_) of the MT ends over each time step was recorded. The displacements and corresponding time steps were then summed over all individuals and over the total measurement period. For the simulation data, the lengths of all individuals are known at all times, so ∑ *t*_i_ = (number of MT seeds) * (total time of measurement). To apply the *J*_TimeStep_ equation to experimental data, it is not necessary for the same individuals to be observable over all time steps; for such data, the sums would include only those displacements that are observed. For additional information about our time-step analysis, please see the Supplemental Methods of (Jonasson et al., 2019).

## Acknowledgments

This work was supported by NSF grants MCB-1244593 and MCB-1817966 to HVG and MCB-1817632 to EMJ.

## Data Availability Statement

The simulation code, analysis codes, and data are available from the corresponding author upon request.

