## SupplementalFiguresS1_S2 for "Relationship between Dynamic Instability of Individual Microtubules and Flux of Subunits into and out of Polymer"

### SUPPLEMENTAL FIGURES

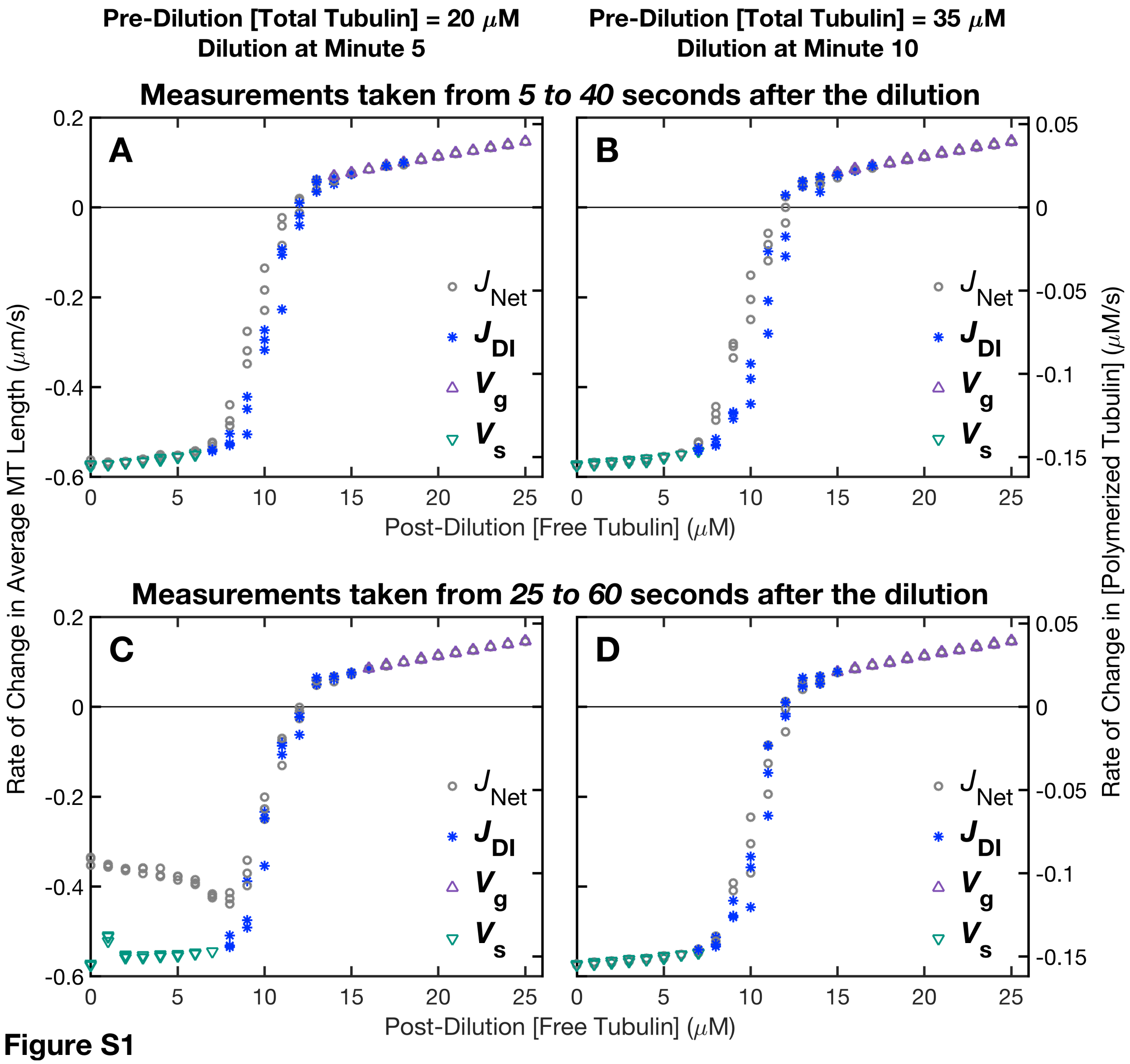


**Figure S1**

**Supplemental Figure S1: Companion figure to Figure 9.** $J_{Net}$ and $J_{DI}$ are re-plotted from **Figure 9**. $V_s$ is plotted when time in growth is zero and $V_g$ is plotted when time in shortening is zero. ***Mathematical rationale***: If time in growth is 0, then $F_{cat}$ is undefined. If time in shortening is 0, then $F_{res}$ is undefined. Therefore, in **Figure 9**, $J_{DI}$ is plotted only when time in growth and time in shortening are both non-zero. In the limit as time in growth goes to 0, $F_{cat}$ goes to infinity, so $F_{res}$ / ($F_{res}$ + $F_{cat}$) approaches 0 and $F_{cat}$ / ($F_{res}$ + $F_{cat}$) approaches 1. Thus, $J_{DI}$ approaches $V_s$. Similarly, as time in shortening goes to 0, $F_{res}$ goes to infinity, so $F_{cat}$ / ($F_{res}$ + $F_{cat}$) approaches 0 and $F_{res}$ / ($F_{res}$ + $F_{cat}$) approaches 1. Thus, $J_{DI}$ approaches $V_g$. ***Methods***: $J_{Net}$ and $J_{DI}$ are re-plotted from **Figure 9**. $V_g$ and $V_s$ were obtained using the DI analysis method (Methods, Section 3.2.2). In panel **A**, the $V_s$ and $V_g$ data are subsets of the $V_s$ and $V_g$ data plotted in **Figure 4A**. ***Interpretations***: $V_g$ and $V_s$ can be used to extend the $J_{DI}$ curve to the ranges where time in growth or time in shortening is zero. The $V_s$ in panel **C** does not shift up as $J$ does. Thus, the shift in $J$ is not due to a shift in $V_s$. Rather, when complete depolymerizations occur, the proportion of time in shortening is not 1, so $J$ does not equal $V_s$.

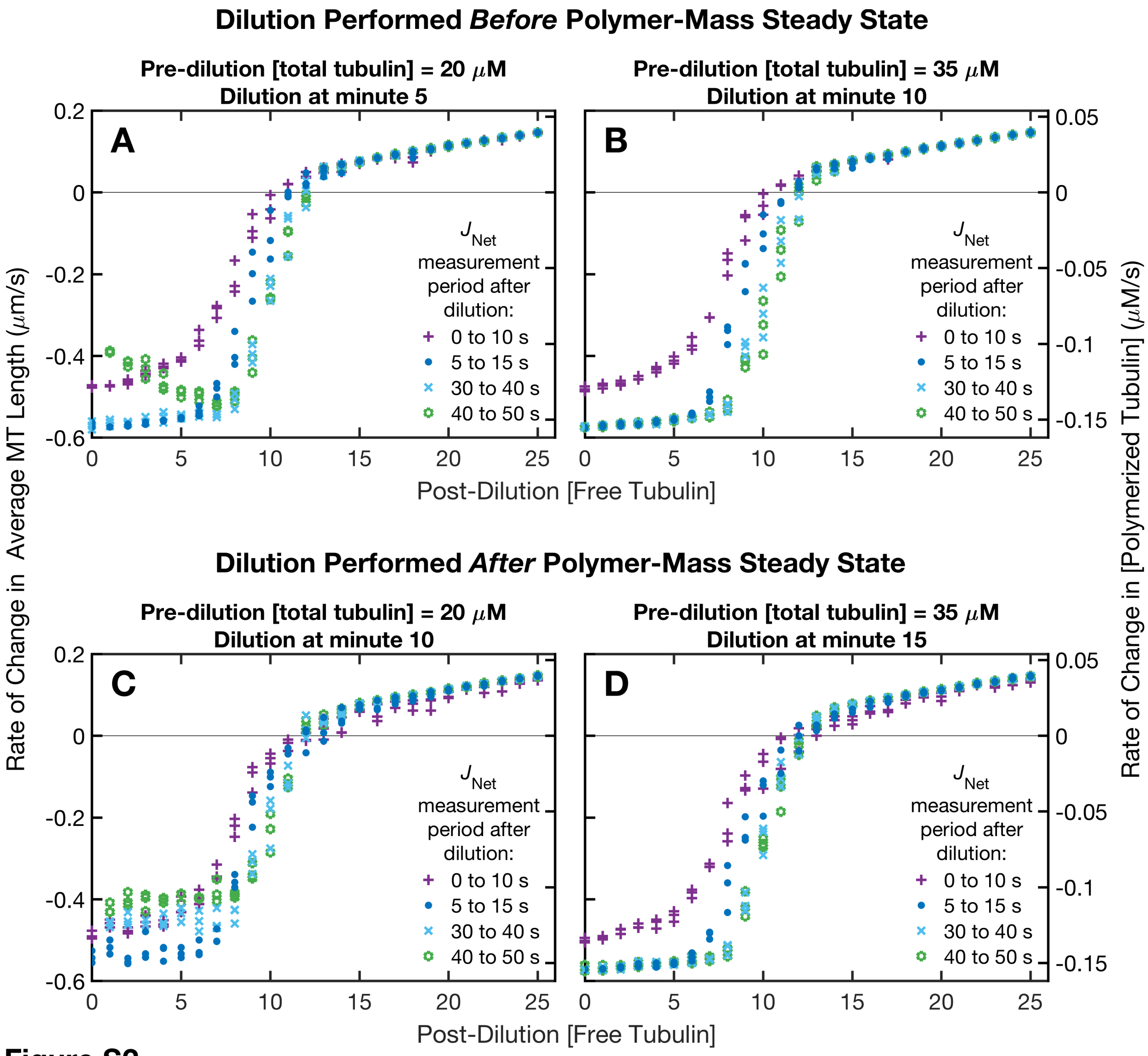


**Figure S2**

**Supplemental Figure S2: Companion figure to Figure 11.** Measurements of $J_{Net}$ are shown for 10-second intervals with varying start times after the dilution, as indicated in the keys on the plots. Panel **D** is re-plotted from **Figure 11**. The dilution was performed either before (panels **A-B**) or soon after (panels **C-D**) polymer-mass steady state (**Table 1**) was reached in the pre-dilution competing systems. ***(A,C)*** Pre-dilution [total tubulin] = 20 µM, with either time of dilution = minute 5 of the simulation (panel **A**), or time of dilution = minute 10 (panel **C**). **(*B,D*)** Pre-dilution [total tubulin] = 35 µM, with either time of dilution = minute 10 (panel **B**), or time of dilution = minute 15 (panel **D**). ***Methods***: Data points are plotted for each of three independent replicates of the simulations at each value of post-dilution [free tubulin]. ***Interpretations***: *Effect of changing the post-dilution start time of the measurement period*: In all panels, a delay after the dilution is needed for *J* to reach it steady-state value for the post-dilution [free tubulin]. As noted in the **Figure 11** legend, the delay is needed to allow the GTP cap (e.g., as quantified by the number of GTP-subunits) to adjust to the new [free tubulin]. However, if the delay is too long (e.g., 40 – 50 s in each of panels **A,C**), then the lower arm of *J* begins to shift upwards, as occurs here in the case of the lower pre-dilution [total tubulin] (panels **A,C**), but not for the higher pre-dilution [total tubulin] (panels **B,D**). As will be examined further in Section 2.6.2 and

**Figures 12-15**, this shift is due to complete MT depolymerizations occurring during the measurement period. *Effect of dilution time relative to polymer-mass steady state*: In the experimental literature (e.g., (Carlier et al., 1984a)), dilutions are often performed after the pre-dilution competing system has reached polymer-mass steady state (panels **C-D**). The results in **Figures 12-15** will provide evidence that the dilution should be performed before polymer-mass steady state (panels **A-B**). However, comparing the top and bottom rows of this figure shows that if the dilution is performed before polymer-mass steady state, then the post-dilution delay before measurement is particularly important for identifying the [free tubulin] at which *J* crosses 0, i.e., $CC_{NetAssembly}$ (**Figure 2** and **Table 1**). More specifically, if there is no post-dilution delay before starting measurements (0 - 10 s data series), then performing the dilution before polymer-mass steady state will cause the *J* curve to be shifted upwards (panels **A-B**) relative to where the *J* curve would be if the dilution were performed after polymer-mass steady state (panels **C-D**). *Explanation*: The observation in the previous sentence can be explained by considering the [free tubulin] and GTP cap size at the time of the dilution. First, as shown in (Jonasson et al., 2019), [free tubulin] is higher before polymer-mass steady state than at polymer-mass steady state. So, [free tubulin] at the time of the dilution will be higher in panels **A-B** (dilution performed before polymer-mass steady state) than in panels **C-D** (dilution performed after reaching polymer-mass steady state). Second, the GTP cap size at the time of dilution will depend on the [free tubulin] at the time of dilution. When [free tubulin] is higher, the GTP cap would be larger. Thus, the cap size would be expected to be larger at the time of the dilution in panels **A-B** than in panels **C-D**, consistent with the above observation that the *J* curve in the 0 - 10 s data series is higher in panels **A-B** than in panels **C-B**.
